# Co-transcriptomic network analysis of host-parasite interactions reveals genes underpinning parasite fitness

**DOI:** 10.64898/2026.10.23.761410

**Authors:** Tyler R. Inskeep, Erick Celis, Anthony Nguyen, Simon C. Groen

**Affiliations:** Department of Nematology, University of California, Riverside, CA, USA 92521; Department of Botany and Plant Sciences, University of California, Riverside, CA, USA 92521; Center for Plant Cell Biology and Institute for Integrative Genome Biology, University of California, Riverside, CA, USA 92521

**Keywords:** systems genetics, host-parasite interaction, root-knot nematode, *Meloidogyne hapla*, *Medicago truncatula*, quantitative disease resistance

## Abstract

Interactions between parasites and their host organisms generate continuous variation in infection outcomes. Although dual transcriptome studies have demonstrated extensive molecular dialogues between interacting species, it is still unclear how natural variation in host and parasite gene regulatory networks and coordination between them contribute to diverse disease phenotypes. Here, we re-contextualized co-transcriptomes from recombinant inbred lines of the plant-parasitic root-knot nematode *Meloidogyne hapla* infecting *Medicago truncatula* host plants with relative measures of ‘infection intensity’ as a proxy for parasite load. We used infection intensity to estimate interaction gradients on gene expression by calculating its covariance with transcript levels. Quantitative analyses of interaction gradients for host and parasite transcriptomes identified major differences in development-related transcriptional programs along the infection continuum. In addition, integrated network analyses connected host cell wall remodeling with parasite protein production and secretion. Silencing of candidate root-knot nematode genes reduced host galling and parasite reproductive fitness, providing experimental support for our network-derived predictions. Together, our findings present an integrated systems genetics framework connecting naturally occurring genetic variation to quantitative disease phenotypes.

## Introduction

Symbiotic interactions between parasites and their hosts produce quantitatively variable infection outcomes. Evolutionary theory predicts that natural genetic variation in either interaction partner can generate such continuous differences in infection outcomes (Thompson 2005; Hanifin et al. 2008). The traits involved in infection emerge from complex molecular dialogues between the organisms upon which selection can act. Across diverse pathosystems, parasite genetic variation has been linked to variation in host immune and physiological states (Wu et al. 2015; Guo et al. 2017; Soltis et al. 2020). While individual genes may exhibit large effects on parasite immunogenicity (Stevens et al. 2024) or host immune activation (Vetter et al. 2012; Winkelmüller et al. 2021), disease is more broadly determined by the collective state of interacting molecular networks during infection (Mine et al. 2018). How natural genetic variation propagates through these networks to generate quantitative variation in infection outcomes remains largely unresolved.

Dual transcriptome profiling of host-parasite interactions provides an unprecedented opportunity to characterize the molecular dialogues of infection through simultaneous quantification of transcriptional responses in both interacting organisms (Westermann et al. 2017). By moving beyond either organism in isolation, co-transcriptomic analyses can reveal coordinated molecular programs that emerge during infection. For example, joint profiling of the sugar beet cyst nematode *Heterodera schachtii* and one of its mustard host plants, the thale cress *Arabidopsis thaliana*, across the nematode life cycle revealed trans-kingdom biosynthesis of vitamin B5, enabling the parasite to exploit its host’s metabolic resources (Siddique et al. 2022). Likewise, cross-species expression quantitative trait locus (eQTL) analysis of Northern root-knot nematode (RKN) *Meloidogyne hapla* parasitism on a legume host, the barrel medic *Medicago truncatula*, demonstrated that naturally occurring parasite genetic variation extensively reshapes host gene expression (Guo et al. 2017). Similar patterns occur in interactions between *A. thaliana* host plants and diverse isolates of the infectious grey mold fungus *Botrytis cinerea*, where parasite genetic variation drives widespread changes in the host transcriptome (Zhang et al. 2019; Soltis et al. 2020). Collectively, these studies demonstrated that genetic variation could reshape transcriptional programs across interacting species. However, they largely focused on genetic mapping or broad transcriptomic comparisons between compatible and incompatible interactions. Co-transcriptomic approaches have identified new loci contributing to disease but thus far provided relatively modest insight into how the dynamics of host and parasite transcriptional networks give rise to continuous differences in infection outcomes.

Plant-parasitic RKNs (*Meloidogyne* spp.) are particularly well suited for mechanistic studies of how genetic variation translates to quantitative phenotypic variation in traits relevant to host-parasite interactions because successful infection requires extensive developmental coordination between organisms. Following invasion of plant roots, second-stage RKN juveniles (J2s) induce formation of multinucleate host cells (‘giant cells’) as feeding sites inside galls through large-scale host transcriptional reprogramming (Barcala et al. 2010; Yamaguchi et al. 2017; Saura-Sanchez et al. 2025; Zadegan et al. 2025), repeated cell-cycle activation (de Almeida Engler et al. 1999; de Almeida Engler et al. 2012), and extensive cell wall remodeling (Bird 1961; Wieczorek et al. 2014; Bozbuga et al. 2018). Simultaneously, the RKNs undergo three successive molts to adulthood (from J3- and J4-stage juveniles to adults). Throughout this process, RKNs depend on sustained secretion of effector proteins for successful infection (Rutter et al. 2022). Consequently, differences in infection outcomes are expected to reflect coordinated changes in the developmental and physiological status of both parasites and hosts.

Here, we leverage naturally occurring genetic variation among recombinant inbred lines (RILs) of the RKN *Mel. hapla* to investigate the molecular basis of quantitative differences in infection outcomes. Leveraging published dual transcriptome data on RKN-infected *Med. truncatula* plants (Guo et al. 2017), we first measure ‘infection intensity’, which we interpret as the relative parasite load of plants infected by each RIL. We quantify infection intensity as the ratio of host and parasite RNA sequencing read mapping rates, finding it captures genetically structured variation in parasite development. Second, building upon quantitative genetic frameworks developed to study continuously varying traits (Lande and Arnold 1983; Hanifin et al. 2008), we estimate interaction differentials and gradients describing covariance between transcriptional programs and infection intensity. Third, we generate an integrated host-parasite co-expression network and identify network modules that vary in activity across the interaction continuum. Finally, we functionally test candidate parasite genes predicted by these analyses, demonstrating that components of the RKN protein production and secretion machinery contribute to parasite fitness through successful infection.

## Methods

### Co-transcriptome alignment

Raw RNA-seq data of *Mel. hapla* RIL-induced galls in *Med. truncatula*, generated by Guo et al. (2017), were downloaded from National Center for Biotechnology Information Sequence Read Archive (BioProject PRJNA229407). Reads were aligned to a concatenated *Med. truncatula-Mel. hapla* reference genome, using HISAT2 v2.2.1 with default parameters (Kim et al. 2019). SAM files were converted to sorted and indexed BAM files using SAMtools v1.22 (Danecek et al. 2021).

Gene-level transcript abundances were quantified from aligned BAM files using featureCounts v2.0.8 (Liao et al. 2014). Reads were assigned to annotated exons using a joint genome annotation (GFF) file. Host and parasite count matrices were separated following quantification and variance-stabilized using the *voom()* function from the limma package in R prior to downstream analyses (Ritchie et al. 2015).

### Infection intensity calculation

Mapping statistics for each sample on a per-contig basis were obtained using the *idxstats* function of SAMtools. Infection intensity (analogous to parasite load) was quantified as:

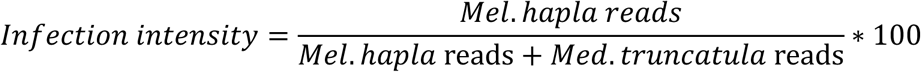

Higher infection intensity values indicate a greater proportion of parasite-derived transcripts within each gall sample. These values were used as the quantitative interaction trait throughout subsequent analyses.

### Variant calling and quantitative genetic analyses

Sequence variants were identified directly from the RNA-seq alignments using the Genome Analysis Toolkit (GATK) v4.2.6.1 (McKenna et al. 2010). Prior to variant calling, read groups were attached to each BAM file and RNA-seq alignments were processed using the *SplicNCigarReads* workflow. Variants were called for each *Mel. hapla* RIL using *HaplotypeCaller* in GVCG mode and jointly genotyped across all samples. Genotype calls were converted into allele dosages (0, 1, or 2), and single nucleotide polymorphisms (SNPs) were filtered based on missing data and minor allele frequency. Following quality control, the resulting matrix of 82,271 SNPs × 80 RILs was used to estimate the additive genomic relationship matrix using the *A.mat()* function implemented in rrBLUP (Endelman 2011).

SNP-based narrow-sense heritability (*h^2^*) was then estimated for infection intensity and individual host and parasite transcript abundances using linear mixed models that included the genomic relationship matrix. Heritability analyses were performed independently for each quantitative trait.

Genome-wide association analyses were performed for infection intensity using GEMMA v0.98.5 (Zhou and Stephens 2012). Association testing was conducted using a linear mixed model that included the genomic relationship matrix to account for relatedness among the *Mel. hapla* RILs. Wald test statistics were used to evaluate marker-trait associations, and genome-wide significance was assessed following Benjamini-Hochberg correction for multiple testing.

### *Mel. hapla* life stage-specific expression inference

To determine whether variation in infection intensity reflected differences in parasite developmental progression, we inferred life stage-specific expression patterns for *Mel. hapla* genes using publicly available developmental transcriptomes from a related RKN species, the congener *Mel. incognita* (Choi et al. 2017). Life stage specificity for each *Mel. incognita* transcript was quantified using a modified version of the tissue specificity statistic *τ* (Yanai et al. 2005), here termed *τ_Life_:*

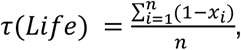

where *x_i_* represents transcript abundance in life stage *i,* normalized to the maximum expression observed across all life stages, and *n* is the number of developmental stages considered.

Orthologous relationships among *Meloidogyne* species were inferred using OrthoFinder (Emms and Kelly 2019). For each *Mel. hapla* gene, all corresponding *Mel. incognita* homologs were identified through one-to-one and one-to-many orthology relationships. Genes with *τ_Life_ > 0.7* were considered developmentally biased and assigned to the life stage exhibiting maximum expression. An inferred ancestral life stage annotation was transferred to *Mel. hapla* only when all corresponding *Mel. incognita* homeologs exhibited concordant life stage-specific expression. This enabled inference of life stage-biased gene expression for 6,262 *Mel. hapla* genes.

To determine whether developmental gene sets contributed to major axes of parasite transcriptional variation, genes assigned to each life stage were tested for enrichment among the 300 transcripts with the most extreme loading scores for each principal component using Fisher’s exact tests. *P*-values were corrected for multiple testing using the Bonferroni method.

### *Med. truncatula* cell type marker gene sets

To determine whether developmentally patterned host transcripts explained variation in infection intensity, publicly available single<u>-</u>nuclei transcriptomes of wild-type *Med. truncatula* accession ‘Jemalong A17’ roots were obtained from Pereira et al. (2024). Only non-nodulated root samples were retained and reanalyzed to identify cell type-specific marker genes. Cell identities were assigned based on the marker genes reported from Pereira et al. (2024).

Marker gene sets for each annotated cell type were then tested for enrichment among the 300 host transcripts with the highest and lowest loading scores for each principal component using Fisher’s exact tests. *P*-values were corrected for multiple testing using the Bonferroni method.

### Interaction differential and gradient estimation

To identify transcriptional programs associated with quantitative variation in infection intensity, we adapted the concepts of selection differentials and gradients from evolutionary quantitative genetics (Lande and Arnold 1983; Hanifin et al. 2008) to transcriptomic traits. Infection intensity was treated as a quantitative interaction trait, and regression coefficients describing covariance between transcript abundance and infection intensity were termed interaction differentials (*S_h_* and *S_p_*). After normalizing infection intensity (dividing by the population average) and standardizing or scaling transcript abundance, interaction differentials were estimated independently for every host and parasite transcript using *glm()* with the formula *infection_intensity ∼ expression* in R. *P*-values were corrected for multiple testing using the Bonferroni method.

To account for extensive correlations among levels of individual transcripts, interaction gradients (*β_h_* and *β_p_*) were inferred for coordinated transcriptional programs. Principal component analyses were performed independently on host and parasite transcriptomes using the *prcomp()* function in R, and sample loading values for PC1-PC11 (host) and PC1-PC10 (parasite) were related to normalized infection intensity using the linear mixed models:

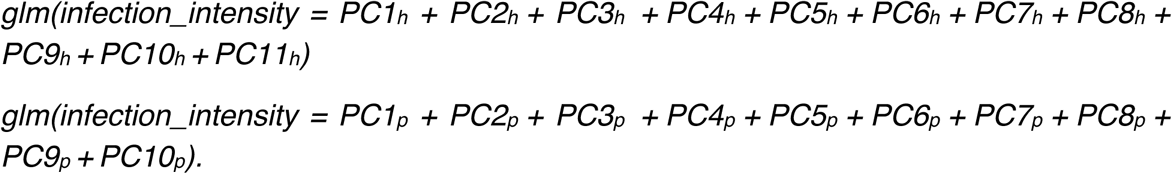

Because principal components are approximately orthogonal, multiple regression coefficients (*β*) describe the direct association between each axis of transcriptional variation and infection intensity (Mitchell-Olds and Shaw 1987).

### Network analysis

Weighted gene co-expression networks were inferred independently for the host and parasite transcriptomes using the WGCNA package in R (Langfelder and Horvath 2008). To reduce noise from weakly varying transcripts, genes were ranked according to their variance across samples prior to network inference. For the *Med. truncatula* transcriptome, the 50% most variable genes (*n* = *9,180*) were retained. For *Mel. hapla*, read depth per sample was comparatively lower, so we applied a more conservative filtering and retained the 25% most variable genes (*n* = *2,159*).

Soft-thresholding powers were selected independently for each organism using the *pickSoftThreshold()* function and an unsigned network topology. The lowest power producing a scale-free topology fit of approximately *R^2^ = 0.8*, while retaining suitable mean connectivity, was selected. Soft-thresholding powers of 6 and 5 were used for the parasite and host networks, respectively. Adjacency and topological overlap matrices were then calculated from pairwise expression correlations, and transcripts were clustered into co-expression modules. Module eigengenes, defined as the first principal component of expression among genes assigned to each module, were used to summarize coordinated module activity across samples.

Selection differentials and gradients were also inferred for host and parasite network module eigengenes using the approaches described above.

To build the co-transcriptome network, we extracted module eigengenes for all modules that retained a significant estimate for *β* (*adj. P < 0.05*) and modeled all pairwise correlations among host and parasite network modules using Spearman rank order correlation. *P-*values were adjusted for multiple testing using the Bonferroni method.

### Gene silencing

Double-stranded RNAs (dsRNAs) targeting *Mh-tmed3* (MhA1_Contig593.frz3.gene3) and *Mh-dpy18c* (MhA1_Contig2702.frz3.gene1) were designed using the IDT RNAi Design Tool. Candidate 21-bp targeting sequences were selected to share no more than 14 continuous bp of identity with any other *Mel. hapla* transcript, as determined by BLAST. Two dsRNA constructs were synthesized for each candidate gene, along with a control dsRNA targeting GFP (Yimer et al. 2023).

*Mel. hapla* strain ‘San Bernardino’ was maintained on *Solanum lycopersicum* cv. ‘Cherry’ tomato plants under greenhouse conditions. RKN egg masses were collected from infected roots by shaking in 10% bleach for 3 min and recovered using stacked 500-mesh sieves. Eggs were purified by flotation in 37% sucrose (5 min, 1,400 × *g*) before hatching for 3 d at 28 °C on a modified Baermann funnel consisting of a sterile 100-µm cell strainer lined with laboratory tissue and suspended over DEPC-treated water.

Approximately 10,000 freshly hatched J2s were incubated in 50 µL of DEPC-treated water containing 5 µg of each of the two dsRNA molecules designed per gene (or 10 µg of the single GFPi dsRNA molecule), 3 mM spermidine, and 50 mM octopamine for 24 h at 26 °C in the dark. Following treatment, 500 J2s were inoculated onto each of 10 two-week-old *S. lycopersicum* cv. ‘Moneymaker’ seedlings grown in 100% blow sand in a greenhouse set to 28 °C under long day conditions (16h light/8h dark).

To validate transcript knockdown, approximately 3,300 dsRNA-treated J2s were retained from each of two biological replicates per gene for RNA extraction. Total RNA was isolated using TRI-zol reagent followed by purification with the Zymo Direct-zol RNA Miniprep Kit according to the manufacturer’s protocol. First-strand cDNA was synthesized using the High-Capacity cDNA Reverse Transcription Kit (Applied Biosystems) and transcript abundance was quantified by RT-qPCR using Luna Universal qPCR Master Mix (New England Biolabs). Relative expression was normalized to succinate dehydrogenase subunit A (SHDA).

Plants were harvested at 21 d after inoculation. Root systems were rinsed to remove sand and stained with 37 mg L^-1^ erioglaucine for 10 min to visualize egg masses, which were counted under a magnifying glass. Following egg mass quantification, roots were cleared in 10% bleach for 5 min, rinsed thoroughly with water, and stained by boiling in acid fuchsin for 2 min to visualize nematodes within the root tissue. Roots were destained by boiling in acidified glycerol (25% glycerol, 0.5% HCl) immediately before counting.

### Data and code availability

Scripts used for all analyses, as well as summary tables (interaction differentials, network modules, etc.) are available on GitHub at: https://github.com/tylerinskeep/mhapla_medicago_interaction_gradients

## Results

### Infection intensity is influenced genetically and captures parasite developmental progression

Guo et al. recently characterized transcriptome variation in root galls of the legume host *Med. truncatula* infected with a RIL population of the RKN parasite *Mel. hapla,* demonstrating extensive parasite genetic control over the host transcriptome (Guo et al. 2017). Despite this genetic control, quantitative measures of infection outcome that can be obtained at comparable scale remain limited, making it difficult to connect parasite genetic variation with phenotypic differences among infections. Recent studies have shown that the relative abundance of host- and parasite-derived transcripts in dual transcriptome datasets can reflect biological variation in parasite load (Westermann et al. 2017; Zhang et al. 2019). Motivated by these observations, we quantified infection intensity as a proxy for parasite load by taking the proportion of mapped reads assigned to the *Mel. hapla* genome for gall samples representing infections by 80 RILs (Figs. 1A-B; Table S1).

**Figure 1.**
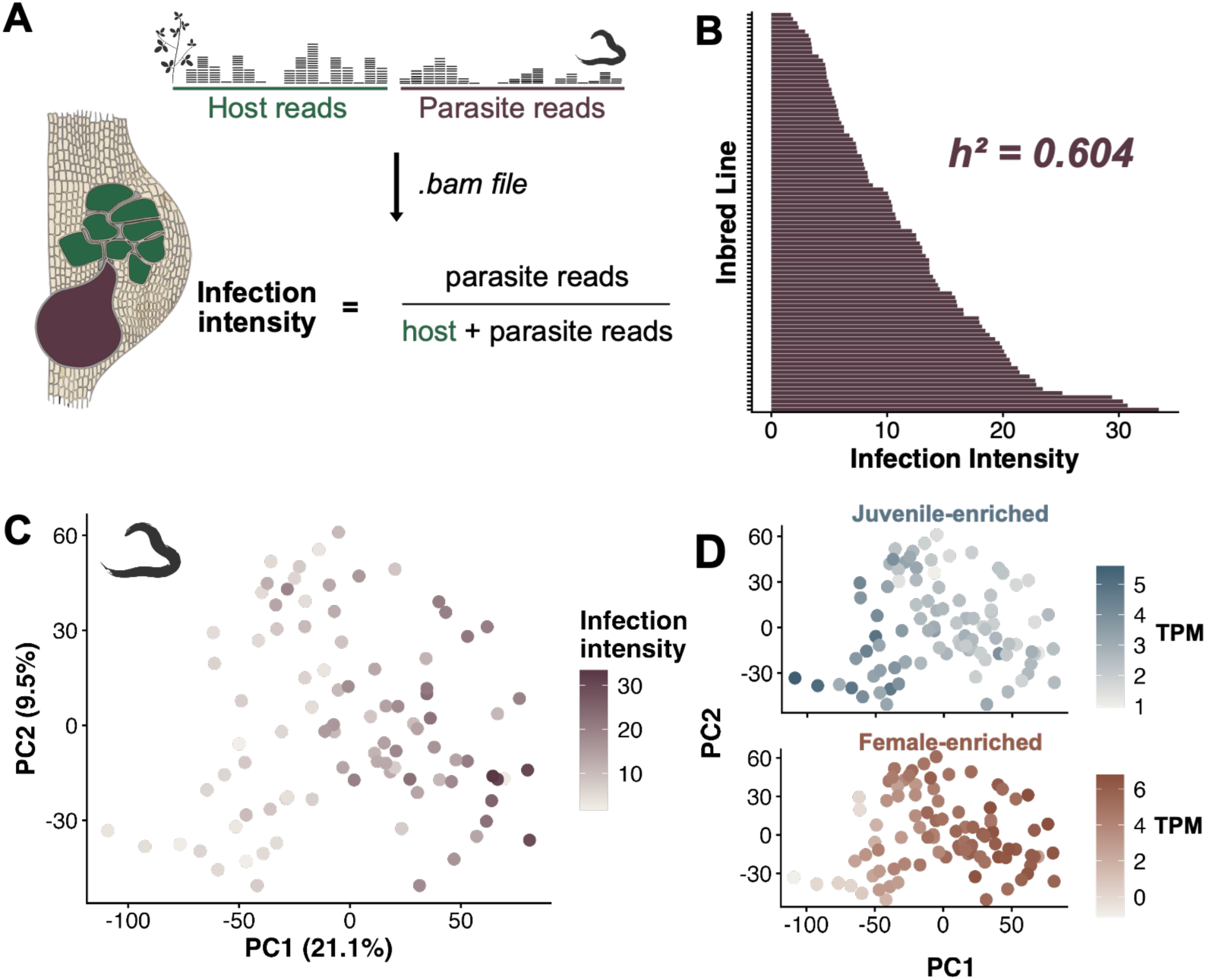
Infection intensity is genetically structured and captures parasite developmental progression. **A.** Infection intensity was quantified for each gall sample as the proportion of mapped RNA-seq reads aligned to the genome of the parasite in this pathosystem, the root-knot nematode *Meloidogyne hapla*, relative to the total number of mapped reads to the genomes of the parasite and its plant host in this system, the legume *Medicago truncatula*. **B.** Infection intensity varied substantially among *Mel. hapla* recombinant inbred lines (RILs) and exhibited high SNP-based narrow-sense heritability (*h^2^*). **C.** Principal component analysis of parasite gene expression revealed that infection intensity varied continuously along the major axes of parasite transcriptome variation. **D.** Expression of the juvenile-enriched transcript *MhA1_Contig2096.frz3.gene8* and the female-enriched transcript *MhA1_Contig298.frz3.gene8* increased towards opposite ends of the transcriptional continuum, indicating that major axes of parasite transcription primarily capture developmental progression.

Several observations indicated that infection intensity reflected biological, and not technical, variation across samples. First, while mapping rates to either genome scaled with library size, infection intensity did not (Fig. S1), implying that it was not an artifact of library size. To determine whether infection intensity was genetically structured, we performed SNP-based genotyping for each RIL using segregating variants identified from the RNA-seq data (Fig. S2A). The resulting kinship matrix was used to estimate SNP-based narrow-sense heritability (*h^2^*; Yang et al. 2017) for infection intensity and for the relative abundance of all host- and parasite-derived transcripts. Infection intensity exhibited substantial heritability (*h^2^ = 0.604*), higher than that observed for expression of most genes (Fig. S2B; Tables S2-3). Despite the strong heritability of infection intensity, no SNP loci explained significant amounts of variation in this trait following Bonferroni correction (*P < 1.4×10^-5^*; Fig. S2C). This pattern is consistent with a genetically complex architecture involving numerous loci of modest effect, which is expected for fitness-proximal traits (Liu et al. 2019; Groen et al. 2020).

Interestingly, we found that infection intensity covaried with the first two principal components (PCs) of *Mel. hapla* gene expression in a nearly continuous trajectory (Fig. 1C). Given the time of sampling (21 days post infection; Guo et al. 2017), we hypothesized that this might reflect quantitative differences in developmental progression towards adulthood across RILs. To address this question, we leveraged developmental transcriptome data from *Mel. hapla*’s triploid congener *Mel. incognita* (Fig. S3A; Choi et al. 2017), and used expression conservation across *Mel. incognita* homeologs to predict life stage-specific expression profiles for homologous *Mel. hapla* genes (Fig. S3B; Tables S4-5). Across the ∼50% of *Mel. hapla* genes (*n = 6,262*) with sufficient homology, we found genes with fourth-stage juvenile (J4)- and adult female-specific expression profiles enriched among the top genes loading into PC1 (Fig. S3E). We additionally observed the expression of J4- and female-specific genes to covary negatively and positively with infection intensity, respectively (Fig. 1D), suggesting the presence of continuous and genetically influenced variation in developmental speed across the RILs.

### Infection intensity is associated with variation in the activity of gall-associated cell types

Despite the strong association between infection intensity and the major axes of parasite transcriptional variation, infection intensity exhibited little separation along the first two principal components of the host (*Med. truncatula*) transcriptome (Fig. 2A). We therefore tested whether infection intensity was associated with more subtle axes of host transcriptional variation by fitting multivariate linear models to the loadings of each principal component. Following adaptations of the Lande-Arnold framework described by Brodie and colleagues (Lande and Arnold 1983; Hanifin et al. 2008), we interpreted the resulting regression coefficients as interaction gradients (*β*), which quantify the relationship between each axis of host transcriptome variation and infection intensity. This analysis identified significant interaction gradients for host PCs 3 and 4 (Figs. 2B & S4C).

**Figure 2.**
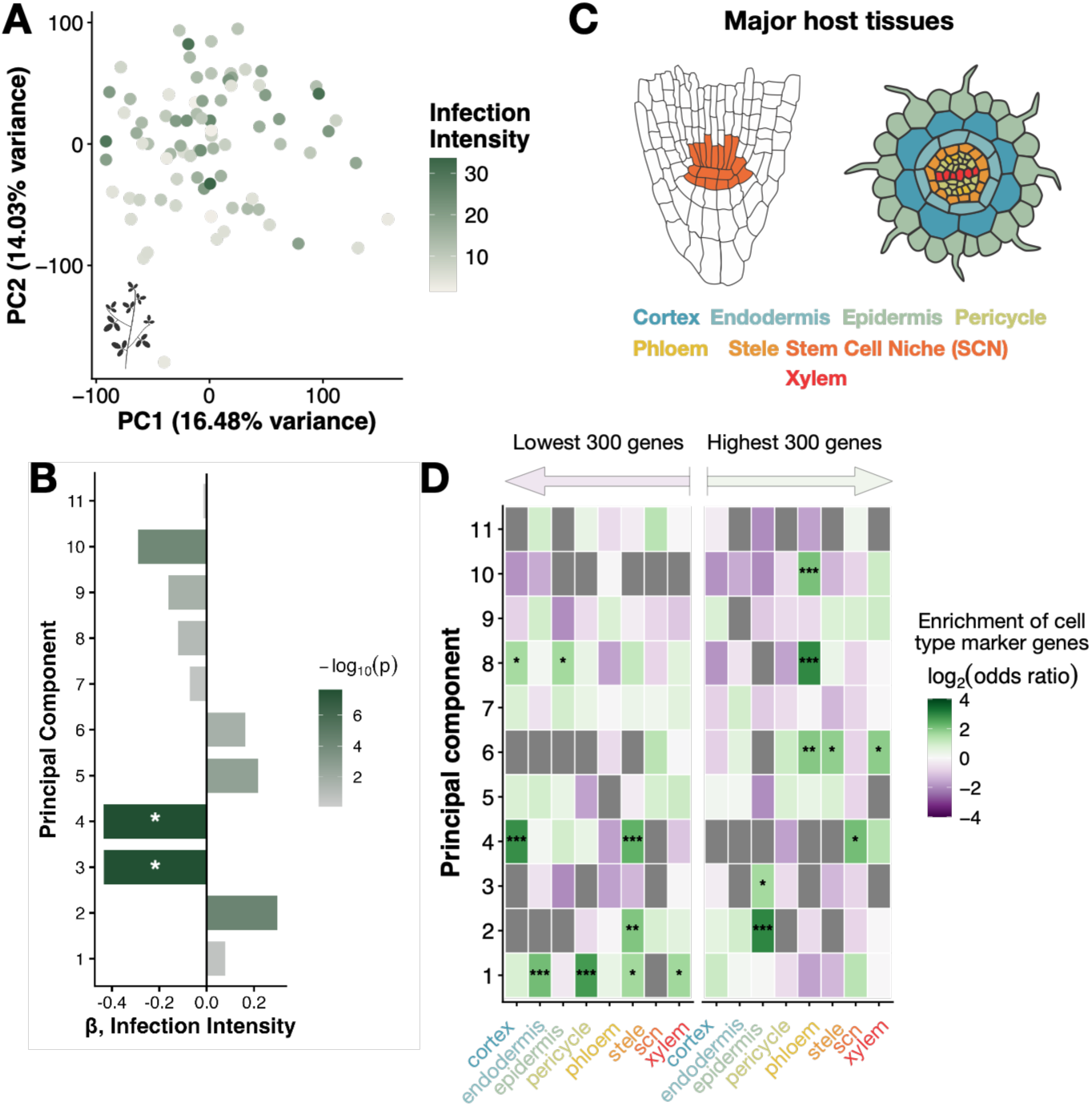
Infection intensity is associated with distinct host transcriptional axes and gall-associated cell types. **A.** Principal component (PC) analysis of *Medicago truncatula* (host) transcriptomes. Samples are colored by infection intensity, which is not associated with the first two axes of transcriptome variation. **B.** Multivariate linear modeling of infection intensity against PC loadings identified significant interaction gradients (*β*) for host PCs 3 and 4. Color denotes statistical significance of *β*. **C.** Schematic of the major root cell types represented by marker gene sets used for cell type enrichment analyses. **D.** Enrichment of cell type-specific marker genes among the 300 genes with the highest and lowest loadings for each PC. PCs 3 and 4 are enriched for epidermal, cortex, stele, and stem cell niche marker genes, indicating that infection intensity is associated with variation in the activity of host tissues undergoing developmental reprogramming during gall formation. Asterisks indicate statistical significance of enrichment test (*\*, adj. P < 0.05*; *\*\*, adj. P < 0.01; ***, adj. P < 0.001*).

To determine the biological processes underlying these interaction gradients, we performed Gene Ontology (GO) enrichment analysis on the 300 genes that loaded most highly into either direction of each principal component. PCs 3 and 4 of host gene expression were enriched for genes involved in DNA replication, mitotic cell cycle progression, microtubule organization, and other processes associated with cell proliferation (Figs.S4D-E). Given that RKN-induced galls are characterized by extensive host cell proliferation during feeding site development (de Almeida Engler et al. 1999; de Almeida Engler et al. 2012), we hypothesized that these interaction gradients reflect differences in the abundance or transcriptional activity of proliferating host cell populations across infections by different *Mel. hapla* RILs.

To address this hypothesis, we retrieved marker genes representing the major cell types of uninfected *Med. truncatula* roots (Pereira et al. 2024; Table S6). Although derived from healthy roots, these provided us with a reference for the cellular origins of transcriptional variation across gall samples. We then tested whether cell type-specific gene sets were enriched among genes with the strongest positive and negative loadings for each principal component. Intriguingly, genes defining PC3 were enriched for epidermal markers, whereas PC4 was enriched for cortex-, stele-, and stem cell niche-specific markers (Fig. 2D), cell types that undergo extensive developmental reprogramming during RKN-induced gall formation (Bird 1961; Yamaguchi et al. 2017; Saura-Sanchez et al. 2025; Zadegan et al. 2025). Altogether, this suggested that infection intensity might have captured relative differences in developmental progression or growth of both host and parasite.

### Interaction differentials identify infection-associated host and parasite transcripts

Having established infection intensity as a biologically meaningful and genetically structured quantitative trait, we next sought to identify the individual host and parasite transcripts associated with variation in infection intensity. To identify such transcripts, we estimated interaction differentials (*S*) for the abundance of every host and parasite transcript, following adaptations of the Lande-Arnold framework by Brodie and colleagues (Lande and Arnold 1983; Hanifin et al. 2008). Interaction differentials quantify the direction and magnitude of covariance between transcript level and infection intensity, with positive values (*S > 0*) indicating transcripts whose expression increases with infection intensity, and negative values (*S < 0*) indicating transcripts whose expression decreases with infection intensity.

Across 8,637 parasite and 18,361 host transcripts, *S* was centered around 0, suggesting that most transcripts exhibit little to no association with infection intensity (Figs. 3A-B; Tables S7-8). Nonetheless, after a conservative Bonferroni correction (*P_p_ < 1.16×10^-6^*; *P_h_ < 5.45×10^-7^*), 1,673 parasite and 357 host transcripts retained significant interaction differential estimates, suggesting infection intensity is associated with changes in a substantial subset of genes in both host and parasite. Gene set enrichment analyses of significant parasite transcripts suggested a major contribution of protein production and secretion to differences in infection intensity, with enrichment of genes involved in translation, the unfolded protein response, and production of the proteinaceous cuticle (Fig. 3C). Additionally, significant host transcripts were enriched for genes involved in biosynthesis and maintenance of the cell wall as well as propagation of stress responses (Fig. 3D).

**Figure 3.**
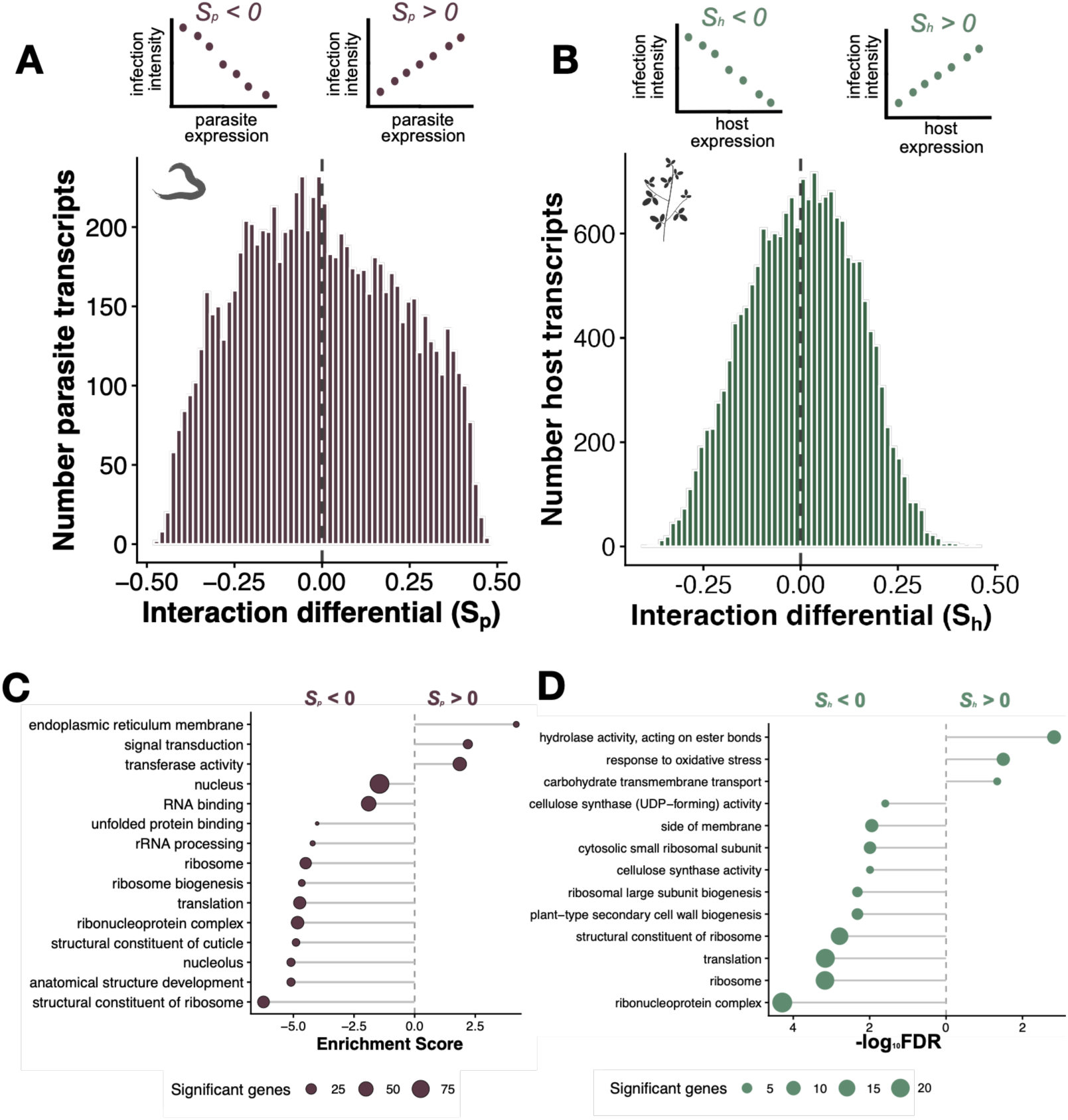
Interaction differentials identify host and parasite transcripts associated with infection intensity. **A-B.** Distribution of interaction differentials (*S*) describing the association between transcript abundance and infection intensity for transcripts of the nematode parasite *Meloidogyne h*apla (*S_p_*; **A**) and plant host *Medicago truncatula* (*S_h_*; **B**). Positive interaction differentials indicate transcripts whose expression increases with infection intensity, whereas negative interaction differentials indicate transcripts whose expression decreases with infection intensity. Insets illustrate the interpretation of positive and negative interaction differentials. **C-D.** Gene Ontology enrichment analysis of transcripts with positive and negative interaction differentials in the parasite (**C**) and host (**D**) transcriptomes.

Gene annotations are relatively limited for *Mel. hapla.* However, given the extensive functional annotation of the *Med. truncatula* genome, we were also able to ask whether any host transcripts with significant estimates of *S_h_* (*S* for host transcripts) have been previously implicated in plant-RKN interactions. Interestingly, the *Med. truncatula* homolog of *12-OXOPHYTODIENOATE (OPDA) REDUCTASE 2* (*MtOPR2*) encoded one of the transcripts with the lowest estimates of *S_h_* (*-0.340, adj. P = 1.6×10^-7^;* Figs. S5A,C). OPR2 is an enzyme involved in oxylipin metabolism and jasmonic acid biosynthesis (Schaller et al. 2000). In *Arabidopsis thaliana*, *AtOPR2* acts as a positive regulator of plant defense to RKNs (Verhoeven et al. 2023). The negative interaction differential observed for *MtOPR2* is therefore consistent with increased expression of a resistance-associated gene in samples with lower infection intensity.

Conversely, we observed strong, positive covariance between the abundance of *TRANSLATIONALLY CONTROLLED TUMOR PROTEIN* (*MtTCTP*) and infection intensity (*S_h_ = 0.3604, adj. P = 6.3×10^-6^*, Figs. S5A-B). *TCTP* homologs encode conserved regulators of cell growth and proliferation in plants (Berkowitz et al. 2008; Brioudes et al. 2010) and a tomato homolog was originally identified as *Meloidogyne-induced giant cell protein DB141* (*MIPDB141*) because of its strong enrichment in RKN-induced giant cells (Jiang et al. 2026). The positive covariance is therefore consistent with our hypothesis of increased gall development in more intense infections.

### Integrated network analyses reveal dynamic and infection-associated host and parasite gene modules

Although our interaction differentials identified individual transcripts associated with infection intensity, they did not capture the coordinated organization of transcriptome variation. We therefore performed parallel gene co-expression network inference of host and parasite transcriptomes to identify gene expression modules associated with infection intensity (Langfelder and Horvath 2008). We identified 9 parasite and 31 host co-expression modules (Tables 1, S9-10). Each module can be represented by a vector, the eigengene, that summarizes the dominant pattern of expression among constituent genes across samples (Tables S11-12). Estimating an interaction differential (*S*) for each module’s eigengene allowed us to look for network modules whose coordinated activity changes dynamically across infection in association with infection intensity. Seven of the 9 parasite modules retained significant estimates of *S_p_* following Bonferroni correction (*P < 0.005*; Figure 3.9B), all of which exhibited dynamic activity along the major parasite transcriptome axes (Fig. S6; Table 1). Additionally, 8 host modules covaried significantly with infection intensity as well (Fig. S7; Table 2). Together, these demonstrate that infection intensity is associated with variation in both host and parasite network states.

**Table 1.** Quantitative genetics of parasite network modules.

| **Module** | **Nr. of Genes** | **$\beta$** | **Significance of $\beta$** | **Eigengene $h^2$** |
| --- | --- | --- | --- | --- |
| parasite1 | 681 | -0.291 | 8.02E-06 | 0.02 |
| parasite2 | 27 | -0.229 | 0.001 | 0.04 |
| parasite3 | 319 | -0.394 | 9.76E-12 | 0.019 |
| parasite4 | 1496 | 0.148 | 0.152 | 0.017 |
| parasite5 | 22 | 0.134 | 0.281 | 0.038 |
| parasite6 | 42 | 0.349 | 9.26E-09 | 0.054 |
| parasite7 | 779 | 0.445 | 2.50E-16 | 0.017 |
| parasite8 | 252 | 0.264 | 9.53E-05 | 0.010 |
| parasite9 | 2860 | 0.241 | 5.88E-04 | 0.112 |

**Table 2.**
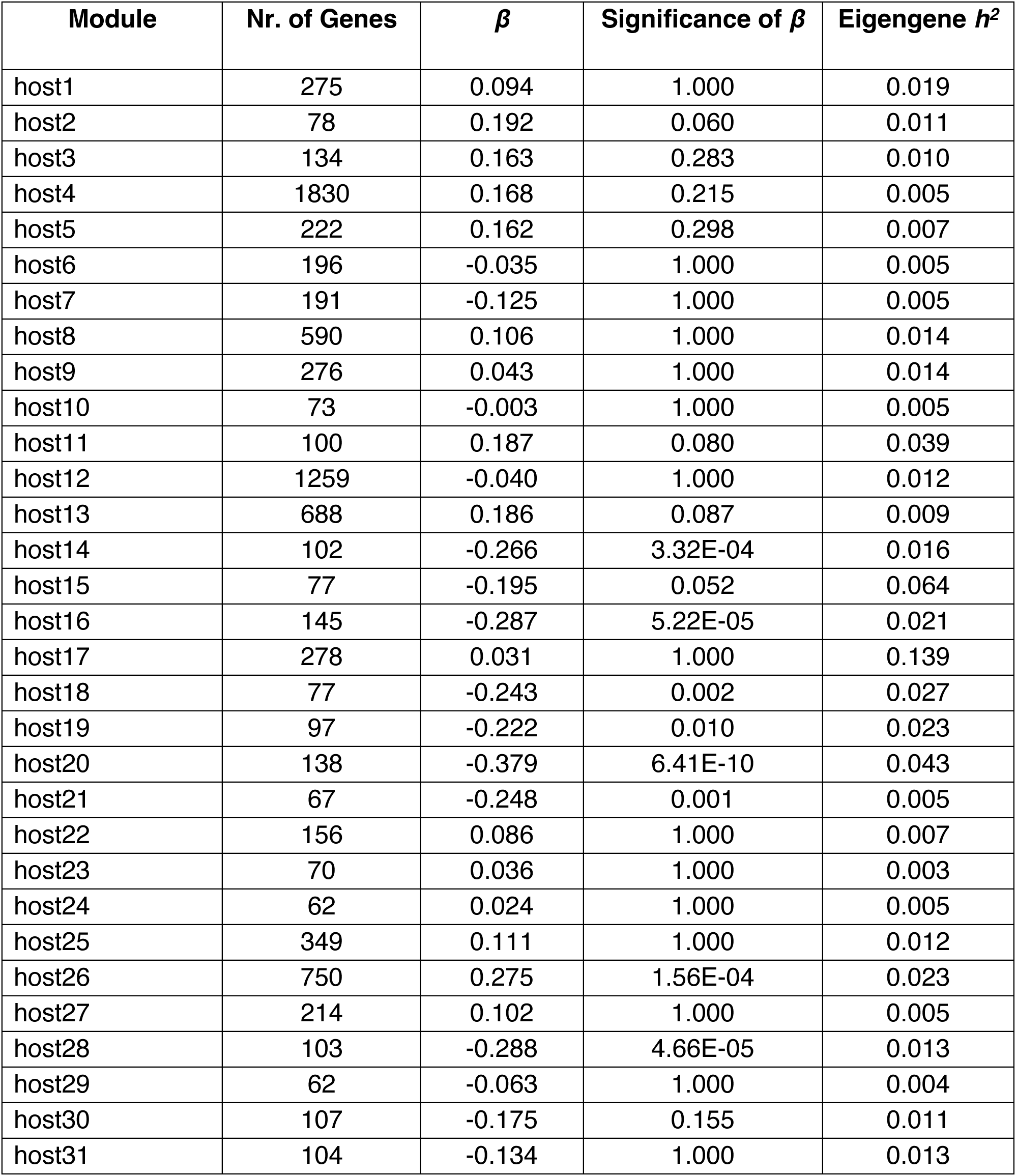
Quantitative genetics of host network modules.

These analyses did not reveal, however, whether host and parasite network states vary independently or as components of a globally coordinated network. To address this question, we next estimated multivariate interaction gradients (*β*; Hanifin et al. 2008) for host and parasite module eigengenes, allowing us to identify modules directly associated with infection intensity, while accounting for covariance among all network states (Tables 1-2). We then quantified pairwise correlations among significant host and parasite modules to reconstruct an integrated host-parasite co-transcriptome in relation to infection intensity (Fig. 4). Five parasite co-expression modules exhibiting significant interaction gradients were correlated with one or more host modules, forming an extensively interconnected network of coordinated transcriptional programs. Indeed, nearly every significant host module was correlated with multiple parasite modules, indicating that the infection continuum is likely a consequence of polygenic network crosstalk.

**Figure 4.**
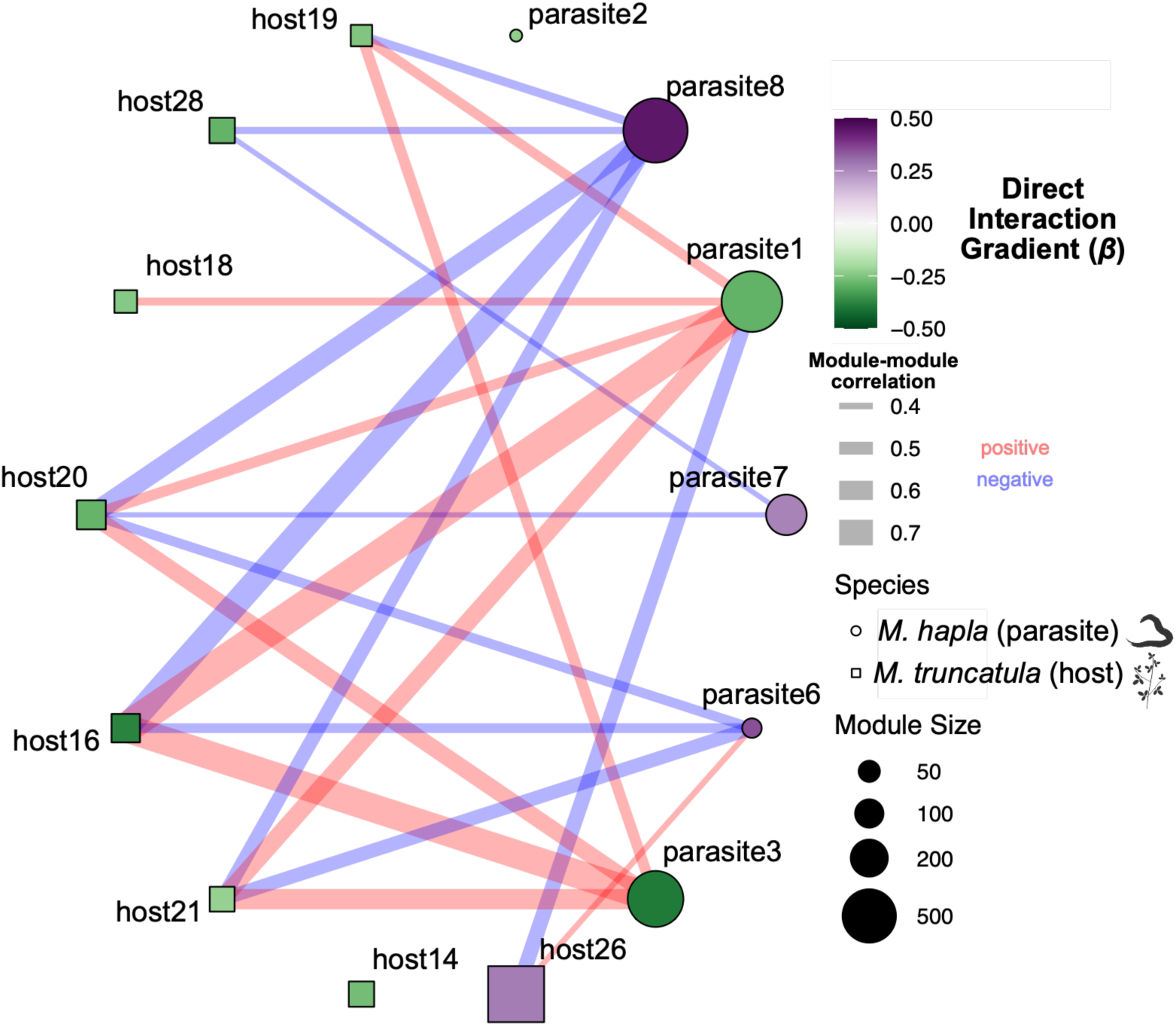
Integrated network analysis reveals coordinated host and parasite interaction gradients. Co-expression modules of the plant host *Medicago truncatula* (squares) and the nematode parasite *Meloidogyne hapla* (circles) exhibiting significant multivariate interaction gradients (*β*) were assembled into an integrated network. Node color represents the standardized interaction gradient (*β*), with positive values indicating modules whose average expression increases with infection intensity and vice versa. Node size is proportional to the number of genes assigned to each module. Edges connect significantly correlated host and parasite module eigengenes, with edge width proportional to the magnitude of the correlation coefficient and edge color indicating positive (red) or negative (blue) correlations. Together, the network illustrates how coordinated transcriptional programs from both organisms jointly define the host-parasite interaction continuum.

### The integrated host-parasite network links host cell wall remodeling with parasite protein production and secretion

To better understand the biological processes represented within the integrated host-parasite network, we examined one of the strongly correlated module pairs. Module ‘host26’ was unique among host modules, being the only host module that positively covaried with infection intensity (*β_h_ = 0.27*; Fig. 4). GO enrichment analysis suggested that ‘host26’ was dominated by genes associated with cell wall remodeling and growth (pectin and lignin catabolism, apoplast, etc.; Fig. 5A). These processes are consistent with the extensive cell wall remodeling required for giant cell expansion, in which pectin-rich cell walls undergo continuous modification during feeding site development (Jammes et al. 2005; Wieczorek et al. 2014; Yamaguchi et al. 2017; Bozbuga et al. 2018). Given this enrichment, we hypothesized that ‘host26’ reflected differences in cellular proliferation and tissue remodeling during gall growth, potentially including proliferation of cortical cells and cell wall modification necessary to accommodate the growing nematode. Consistent with this hypothesis, the module was enriched for previously identified *Med. truncatula* stele marker genes (Odds Ratio = 2.46, adj. *P =* 4.36×10^-3^; Fig. 5B; Pereira et al. 2024), including two encoding pectate lyase enzymes (*MedtrA17_06g0458201, MedtrA17_01g0169731*).

**Figure 5.**
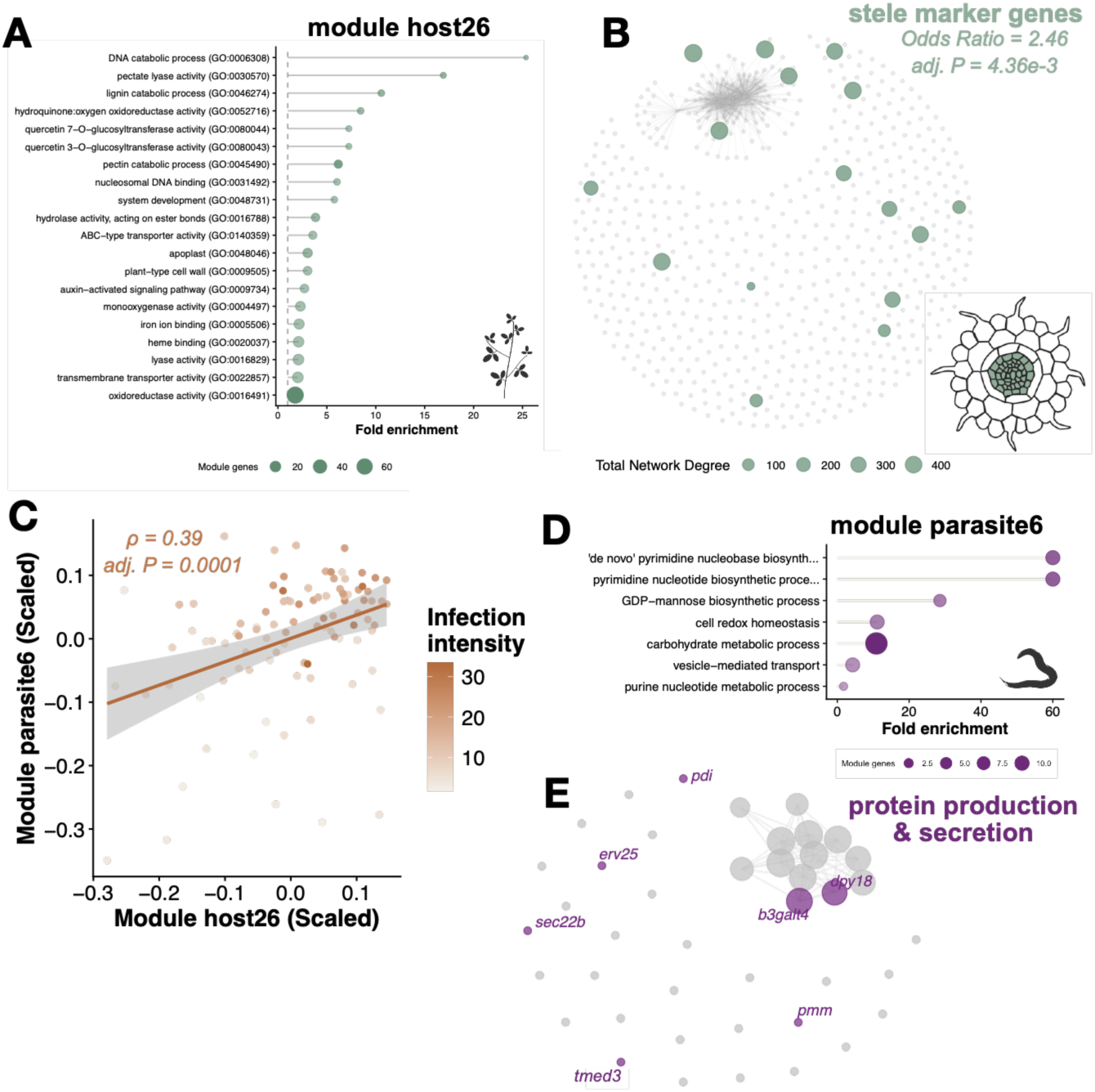
Integrated host and parasite network modules implicate coordinated host cell wall remodeling and parasite protein production during infection. **A.** Gene Ontology (GO) enrichment of co-expression module ‘host26’ of the plant *Medicago truncatula*. Enriched biological processes mostly involve cell wall metabolism. Point size corresponds to the number of module genes assigned to each GO term. **B.** Gene regulatory network (GRN) representation of ‘host26’, enriched for previously determined stele-specific marker genes. Each dot corresponds to one transcript, stele-enriched transcripts are colored green, and their size is proportional to their degree (number of connections) in ‘host26’. **C.** Eigengenes for ‘host26’ and the module ‘parasite6’ of the nematode parasite *Meloidogyne hapla* correlated positively (*ρ = 0.39, adj. P = 0.0001*). Each dot represents a sample, colored by infection intensity, highlighting positive correlations for each module with infection intensity. **D.** GO enrichment of parasite co-expression module ‘parasite6’. Enriched biological processes are associated with protein production and secretion. **E.** GRN representation of ‘parasite6’, highlighting several genes involved in protein secretion in purple.

The activity of module ‘host26’ was positively correlated with that of module ‘parasite6’ (Spearman *ρ = 0.39; adj. P = 0.001*; Fig. 5C), and this parasite module also exhibited a positive interaction gradient (*β_p_ = 0.35*), suggesting that their coordinated activity might be consequential for infection progression.

Module ‘parasite6’ was enriched for genes associated with nucleotide biosynthesis, carbohydrate metabolism, vesicle-mediated transport, and cellular redox homeostasis (Fig. 5D). Enrichment for several of these processes was driven by shared genes involved in protein glycosylation, protein folding in the endoplasmic reticulum (ER), ER protein export, and collagen maturation (Fig. 5E). Notably, this mirrored the original enrichment of transcripts encoding cuticle-related and ER-localized proteins in the interaction differential models for parasite gene expression (*S_p_*; Fig. 3C), suggesting that remodeling of host cell walls might be functionally or genetically coupled with parasite protein production and secretion.

### Parasite protein production and secretion contribute to successful infection

Having identified this coordinated host-parasite transcriptional module, we next asked whether genes associated with parasite protein production and secretion might contribute functionally to quantitative variation in infection success. To address this question, we selected three candidate genes that provided complementary evidence for contribution to differences in infection intensity.

We first focused on *Mh-tmed3* (MhA1_Contig593.frz3.gene3), encoding a member of the broadly conserved p24 family of transmembrane proteins that regulate the vesicular transport of folded proteins from the ER to the Golgi apparatus (Springer et al. 2000). Notably, *Mh-tmed3* was a member of ‘parasite6’, the parasite co-expression module associated with host module ‘host26’ as described above (Fig. 5E). *Mh-tmed3* exhibited a significant interaction differential (*S_p, Mh-tmed3_ = 0.324, adj. P = 0.012;* Fig. S8A). Although this gene lacked expression divergence among the parental isolates, transgressive variation in expression was apparent among the RILs (*P = 0.943*; Fig. S8B, S10; Table S13).

Additionally, we were interested in exploring contributions of *Mh-dpy18c* (MhA1_Contig2702.frz3.gene1) to parasitism. *Mh-dpy18c* encodes a prolyl 4-hydroxylase whose primary substrate is collagen, a key structural component of the nematode cuticle (Hill et al. 2000). Despite lacking a significant interaction differential following multiple testing correction (*S_p, Mh-dpy18_ = 0.29, adj. P = 0.102*; Fig. S9A), *Mh-dpy18c* was a hub gene for module ‘parasite6’ (Fig. 5E) and exhibited expression divergence between the parental strains (*adj. P = 1.6×10^-10^*; Fig. S9B, S10; Table S13). We thus reasoned that *Mh-dpy18c* might contribute to infection outcomes due to its role as a central member of module ‘parasite6’.

To determine whether these genes contribute to parasite performance, we designed and synthesized two double-stranded RNAs (dsRNAs) targeting each candidate gene (Figs. S8D & S9D) along with control dsRNAs targeting the GFP coding sequence, which has no detectable homology to any *Mel. hapla* gene model. Freshly hatched RKN J2s were soaked in a dsRNA-containing solution for 24 hours and used as inoculum for tomato infection assays. Knockdown was assessed via RT-qPCR of the remaining RKNs. Transcript knockdown was successful for both *Mh-tmed3* and *Mh-dpy18c* (Fig. 5A), which motivated us to assess disease-related phenotypes for plants infected with the *TMED3i* and *DPY18ci* RKNs at 21 days post inoculation. Strikingly, transcript knockdown was accompanied by a reduction in host gall formation (Fig. 5B). Furthermore, nematodes inside *TMED3i* and *DPY18ci* galls exhibited delayed development relative to the control, with more juveniles and fewer gravid females depositing egg masses (Figs. 5C-D). Together, these experiments provide functional support for our network analyses, demonstrating that multiple components of the RKN protein production and secretion machinery contribute to successful host plant colonization and reproduction.

## Discussion

Understanding how naturally occurring genetic variation gives rise to continuous differences in infection success remains a central challenge in host-parasite biology. By integrating quantitative genetic analyses with host-parasite co-transcriptomes, we examined how genetically structured transcriptional variation across both host and parasite relates to variation in infection phenotypes. Our analyses revealed numerous interconnected transcriptional modules across both organisms whose expression strongly covaried with infection intensity as a measure of parasite load (Fig. 4). These programs linked host plant pathways implicated in RKN feeding site formation with pathways in the nematode parasite that were associated with protein production and secretion as well as molting, suggesting that successful parasitism emerges from coordinated developmental states across species. Together, these findings support a systems genetics view of host-parasite interactions in which quantitative disease outcomes arise from network-level molecular and cellular processes across the symbiotic organisms.

We demonstrate that infection intensity provides a biologically meaningful and quantitative measure of the outcomes of host-parasite interactions. Infection intensity was strongly influenced by RKN genotype (Fig. S2) and tracked continuous variation in the developmental progression of both host plant and nematode parasite (Figs. 1D, 2D), indicating that this metric reflects biological differences germane to parasite performance rather than stochastic variation in sequencing depth or tissue sampling (Fig. S1). Our findings are consistent with those of previous studies demonstrating that transcript abundance can provide an accurate proxy for microbial biomass or activity within host tissues (Zhang et al. 2019; Akmakjian et al. 2025), as well as recent qPCR-based approaches for high-throughput quantification of RKN infection (Deveux et al. 2025). Thus, by capturing continuous variation in parasite load, infection intensity provided a quantitative phenotype that could be integrated with systems genetics approaches to resolve the underlying network dynamics driving successful infection.

For RKNs as sedentary endoparasites, successful development *in planta* depends on the establishment and maintenance of functional host-derived giant cells that provide nutrition. Accordingly, we reasoned that infection intensity should reflect coordinated variation in both parasite development and host feeding site formation. Consistent with this hypothesis, infection intensity covaried continuously with transcript abundance of host plant genes involved in cell proliferation and cell wall remodeling (Figs. 3D, S4D-E). This was particularly the case for genes enriched in the stele, a tissue that undergoes extensive developmental reprogramming during giant cell formation (Figs. 5A-B; de Almeida Engler et al. 1999; de Almeida Engler et al. 2012; Yamaguchi et al. 2017; Saura-Sanchez et al. 2025; Zadegan et al. 2025). These patterns likely reflect multiple aspects of gall development, including reprogramming of stele-derived cells during giant cell formation, proliferation of surrounding cortical cells, and remodeling of the cell wall to support the growing nematode. These observations were reinforced by transcript-level analyses of interaction differentials (*S_h_*), which identified numerous host genes covarying with infection intensity, including previously characterized regulators of giant cell development and defense against RKNs (Fig. S5). Collectively, these findings support the interpretation that higher infection success emerges through coordinated RKN manipulation of host developmental programs and crosstalk between development and immune signaling.

Within the RKN parasite, pathways underpinning protein secretion and cuticle maintenance repeatedly stood out as covarying with infection intensity (Figs. 5D-E, S8-9). Silencing of *Mh-tmed3* and *Mh-dpy18c* expression significantly reduced RKN fitness on tomato (Fig. 6), providing experimental support that these pathways contribute to successful parasitism. Perturbation of core cellular processes, including proteolysis, can impair development and reproductive fitness in RKNs (Antonino de Souza Júnior et al. 2013), and protein trafficking may be driving infection and fitness of nematode parasites through at least two mechanisms. A first mechanism is that RKN infection is, for an important part driven, by secretion of effector proteins (Rutter et al. 2022), for which core components of the protein production and secretion machinery are key. Sustained delivery of secreted effector proteins into the host plant is necessary throughout the prolonged parasitic phase, suggesting that variation in the cellular capacity to synthesize, process, and traffic secreted proteins may contribute as much to higher infection success as variation in the parasite’s effectors themselves. This interpretation is consistent with observations across diverse host-associated parasites. For example, *ERP3*, a fungal homolog of *TMED3*, is required for full virulence in *Candida albicans* (Yu et al. 2026), while disruption of the *N*-linked glycosylation pathway impairs pathogenicity in the rice blast fungus *Magnaporthe oryzae* by disrupting glycosylation of secreted effector proteins (Chen et al. 2014).

**Figure 6.**
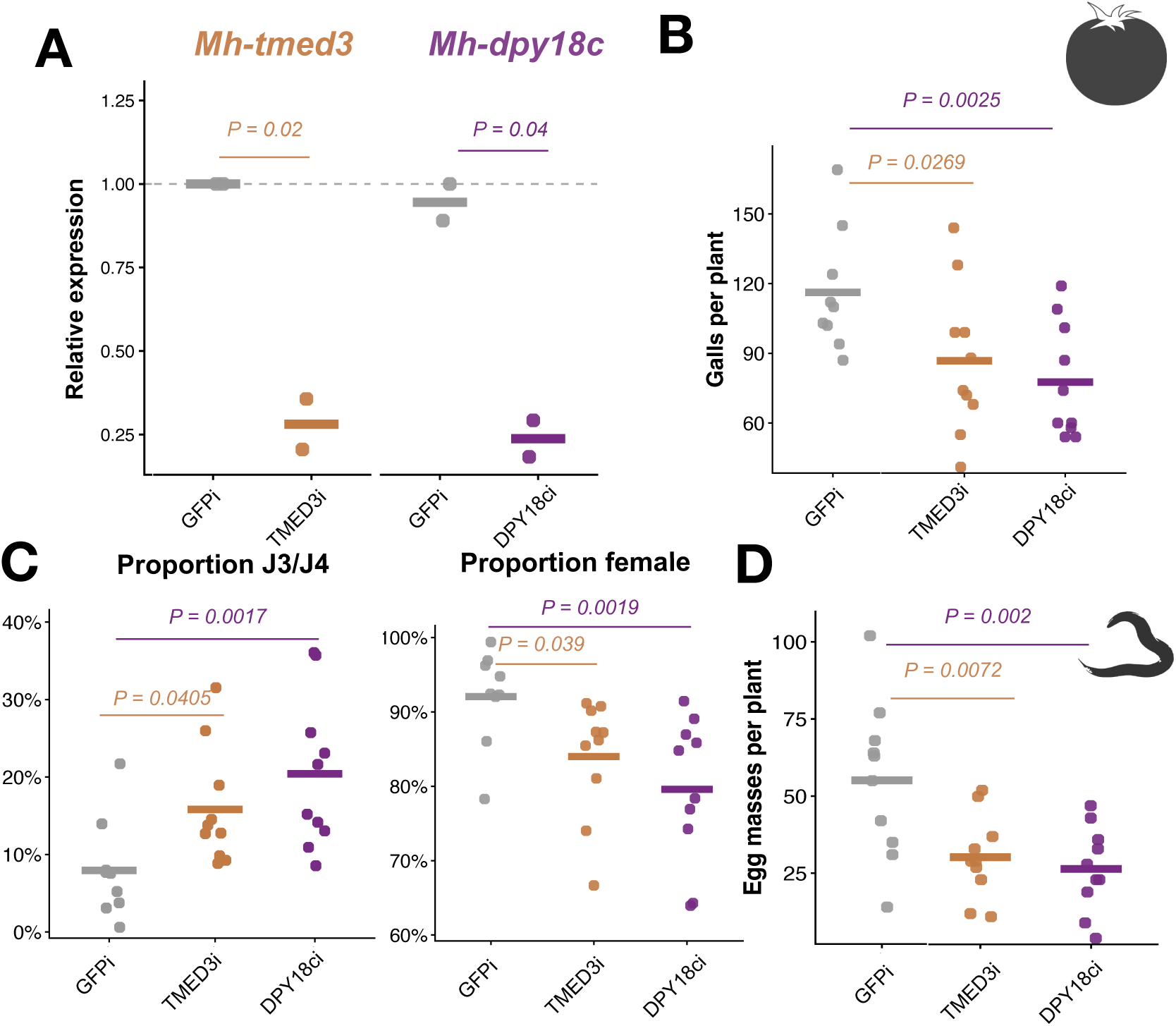
Silencing of candidate parasite genes suppresses parasite fitness. **A.** Relative transcript abundances of *Mh-tmed3*, and *Mh-dpy18c* in the nematode parasite *Meloidogyne hapla*, following 24 hours of dsRNA soaking. Expression was quantified by RT-qPCR and normalized relative to a *GFPi* control (*n = 2*). Horizontal bars indicate treatment means. One-tailed Student’s *t*-tests were used to test for reduced expression in dsRNA-treated nematodes relative to control nematodes. **B.** Galls produced per tomato plant at 21 days post inoculation with dsRNA-treated, second-stage juvenile (J2) nematodes (*n = 10*). **C.** Proportion of nematodes per root system at each developmental stage (*n = 10*). **D.** Egg masses produced per root system (*n = 10*). Counts were analyzed using a negative binomial generalized linear model, with pairwise comparisons performed relative to *GFPi*.

A second mechanism is that protein trafficking is essential for cuticle maintenance, a process fundamental to nematode growth and development. Indeed, multiple lines of evidence implicated cuticle biogenesis and remodeling as determinants of higher infection success. The nematode cuticle undergoes repeated synthesis and remodeling throughout post-embryonic development to provide structural support as nematodes progress through various molts (Page and Johnstone 2007). Consistent with this biology, *Mh-dpy18c*, which encodes a collagen-modifying enzyme critical for cuticle integrity in *Caenorhabditis elegans* (Hill et al. 2000), emerged as a hub gene within module ‘parasite6’ despite only modest transcript-level covariance with infection intensity (*S_p_*; Figs. 5E & S9A).

Beyond its structural role, the cuticle represents a dynamic interface between invertebrate parasites and their hosts. The cuticle surface coat is composed largely of glycoproteins (Robertson et al. 1989) and its continual shedding and renewal throughout host invasion and RKN development is influenced by environmental and host-derived signals (Rogers and Bird 1965; López de Mendoza et al. 2000; Akhkha et al. 2002). In parallel, the biochemical composition of cuticle collagens changes throughout development, modifying the surface chemistry and mechanical properties of the RKN (Reddigari et al. 1986). These changes are thought to protect the parasite from host defenses and natural enemies (Davies et al. 2008; Davies and Curtis 2011). Consistent with this hypothesis, in cyst nematodes, which are relatively closely related to RKNs, peroxidase effectors localize to the cuticle to detoxify host-derived reactive oxygen species (Waetzig et al. 1999). Finally, RNAi-mediated silencing of cuticle collagen genes in *Mel. incognita* reduces female fecundity on tomato (Banerjee et al. 2017), consistent with the reduced egg mass production observed following *Mh-dpy18c* silencing we report here (Fig. 6). Together, these findings suggest that naturally occurring genetic variation in cuticle remodeling may influence infection success by coordinating parasite development, protection, and prolonged host colonization. Future studies addressing the relative contributions of effector protein secretion versus cuticle protein secretion to RKN infection might help further identify novel regulators of virulence.

### Conclusion

Collectively, our findings support a view of quantitative disease outcomes as an emergent property of coordinated developmental programs spanning both host and parasite. Our analyses identified conserved cellular processes involved in protein production and secretion as well as cuticle remodeling in RKNs that likely underpin successful parasitism, potentially independent of individual effectors. Integrating quantitative genetics approaches with our co-transcriptome network allowed us to connect parasite genetic variation to complex disease phenotypes. Because these approaches are broadly applicable across host-parasite systems, they may facilitate the discovery of conserved biological processes underlying quantitative variation in disease outcomes in diverse pathosystems.

## Acknowledgments

We are grateful to members of the Groen lab for helpful discussions. The functional validation experiments would not have been possible without the generous contribution of *Mel. hapla* inoculum by Bao-Lam Huynh and Tra Duong. OpenAI Codex, using the GPT-5.6 Sol model, was used to organize and debug portions of the analysis code. Conceptualization and supervision: T.R.I. and S.C.G.; Data analysis: T.R.I.; Funding acquisition: T.R.I. and S.C.G.; Investigation: T.R.I., E.C., and A.N.; Writing: T.R.I. and S.C.G.

---

**Figure S1.**
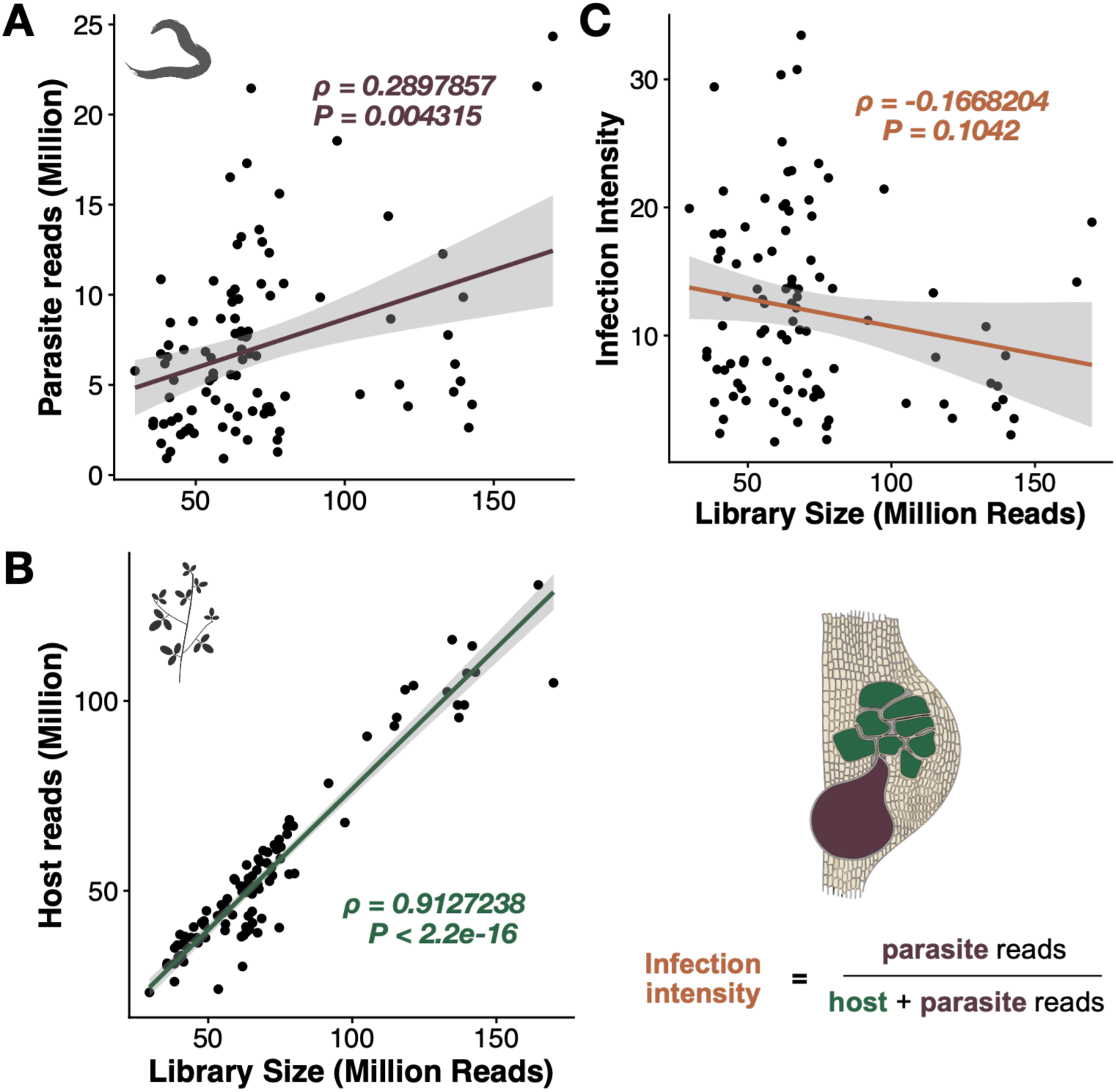
Infection intensity is independent of sequencing depth. **A.** Relationship between total library size and the number of reads mapping to the genome of the nematode parasite *Meloidogyne hapla*. **B.** Relationship between total library size and the number of reads mapping to the genome of the plant host *Medicago truncatula*. **C.** Infection intensity is independent of library size. Lines represent linear regressions with 95% confidence intervals; Spearman’s rank correlation coefficients (*ρ*) and corresponding *P*-values are reported.

**Figure S2.**
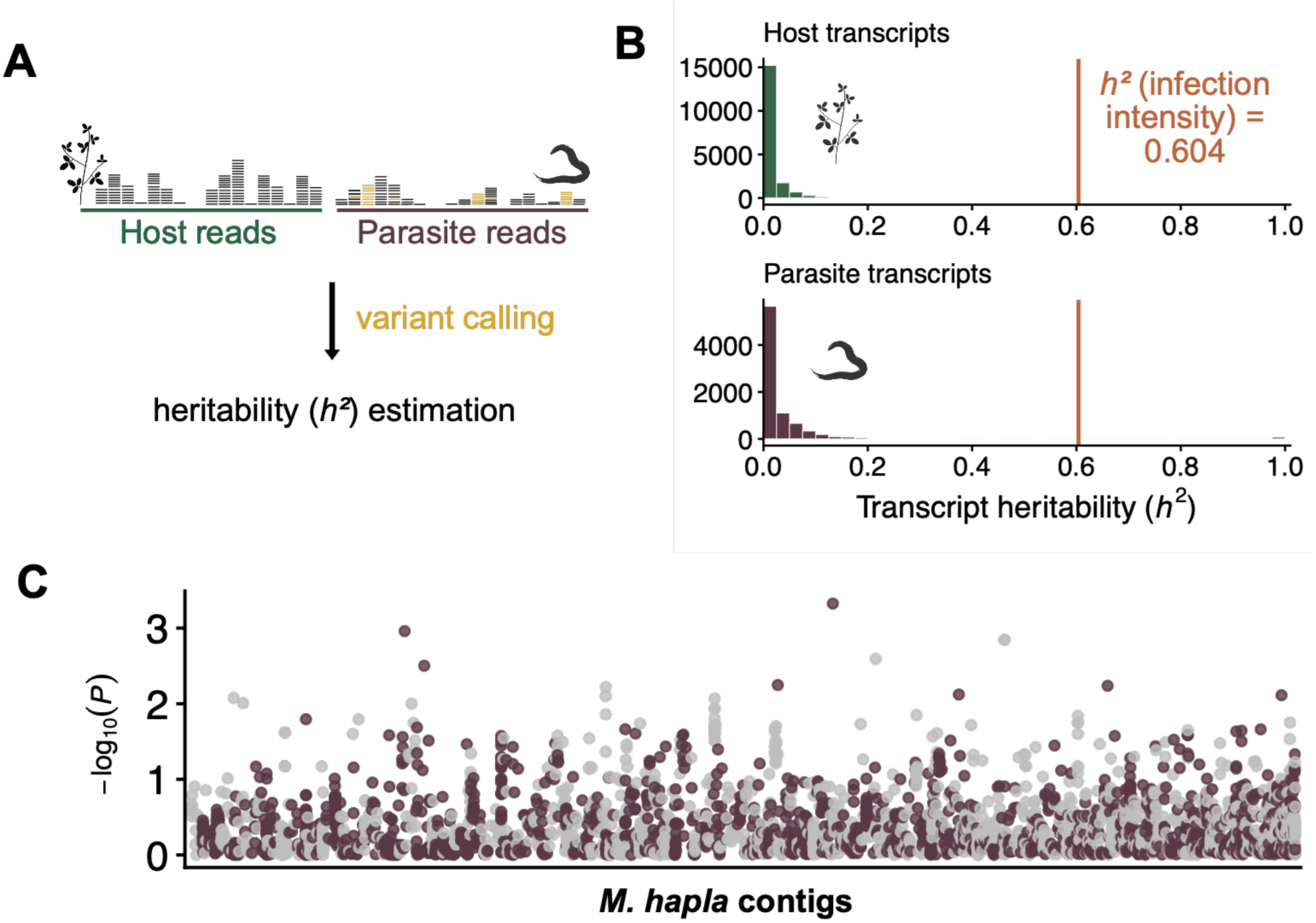
Infection intensity estimates resemble a polygenic trait. **A.** Schematic illustrating SNP-based genotyping of recombinant inbred lines (RILs) of the nematode parasite *Meloidogyne hapla* from RNA-seq data for estimation of narrow-sense heritability (*h^2^*). **B.** Distribution of *h^2^*estimates for abundances of transcripts derived from the plant host *Medicago truncatula* (above) and the parasite (below), *h^2^* for infection intensity (0.604) is indicated by the vertical orange line. **C.** Genome-wide association analysis of infection intensity across *Mel. hapla* contigs. No loci exceeded the genome-wide significance threshold, consistent with a genetically complex architecture.

**Figure S3.**
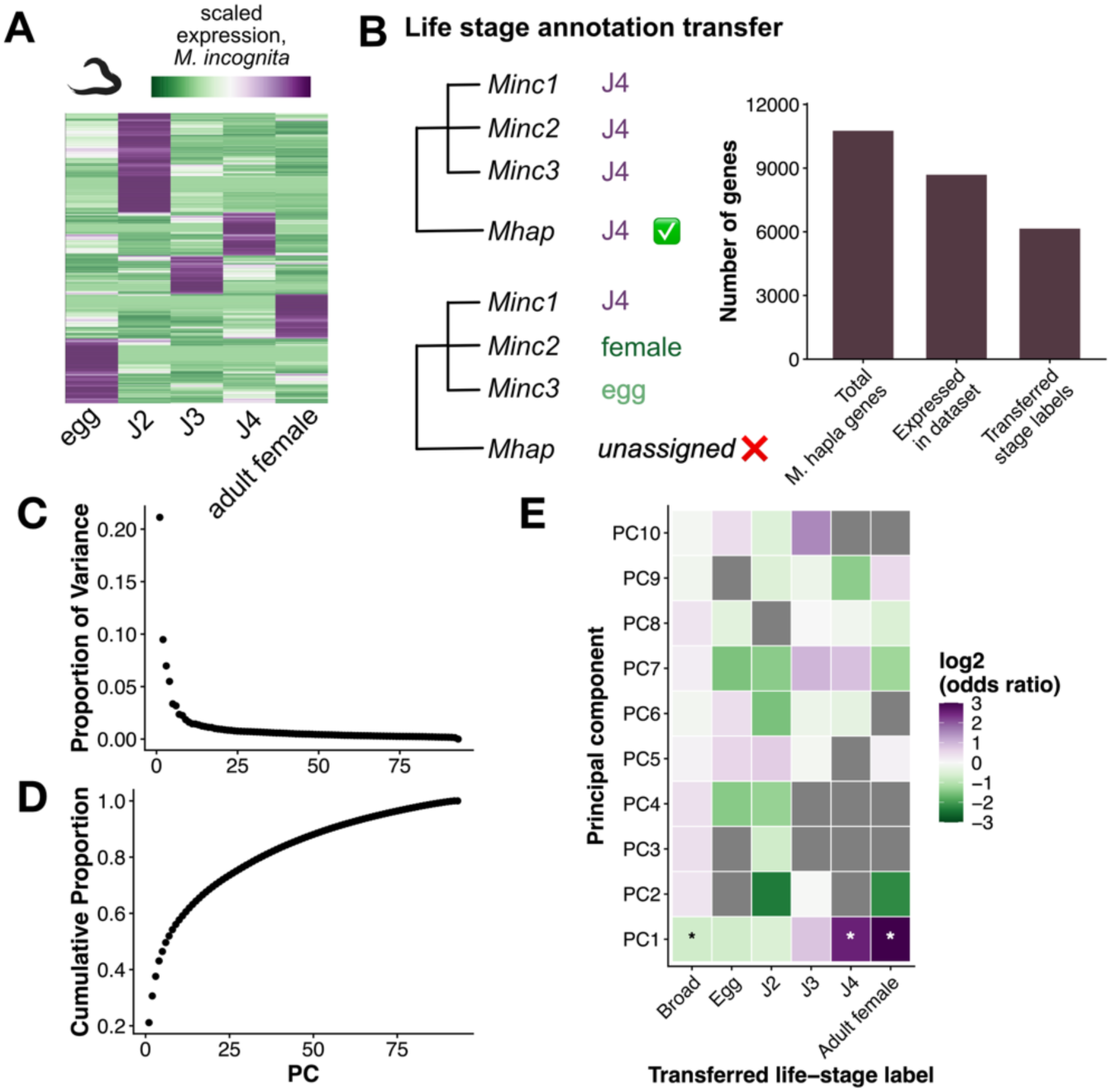
Conserved developmental expression identifies life stage-specific correlates of parasite transcriptome variation. **A.** Developmental expression profiles for genes of the nematode parasite *Meloidogyne incognita* across five parasite life stages—egg; second-, third-, and fourth-stage juvenile (J2, J3, and J4); and adult female—were used to infer stage-specific expression patterns. **B.** Schematic illustrating transfer of developmental stage annotations from *Mel. incognita* (3n) homeologs to orthologous *Mel. hapla* (2n) genes. Stage labels were assigned only when all *Mel. incognita* homeologs exhibited concordant life stage-specific expression. Bar plot summarizes the number of *Mel. hapla* genes in the genome, those detected in the dataset, and those assigned developmental stage annotations. **C.** Variance explained by each principal component of the *Mel. hapla* transcriptome. **D.** Enrichment of inferred life stage-specific genes among the 300 genes with the most strongly positive and negative loadings for each principal component. Tiles with an asterisk represent those with significant enrichment scores, *adj. P < 0.05*. PC1 is enriched for J4- and adult female-specific transcripts.

**Figure S4.**
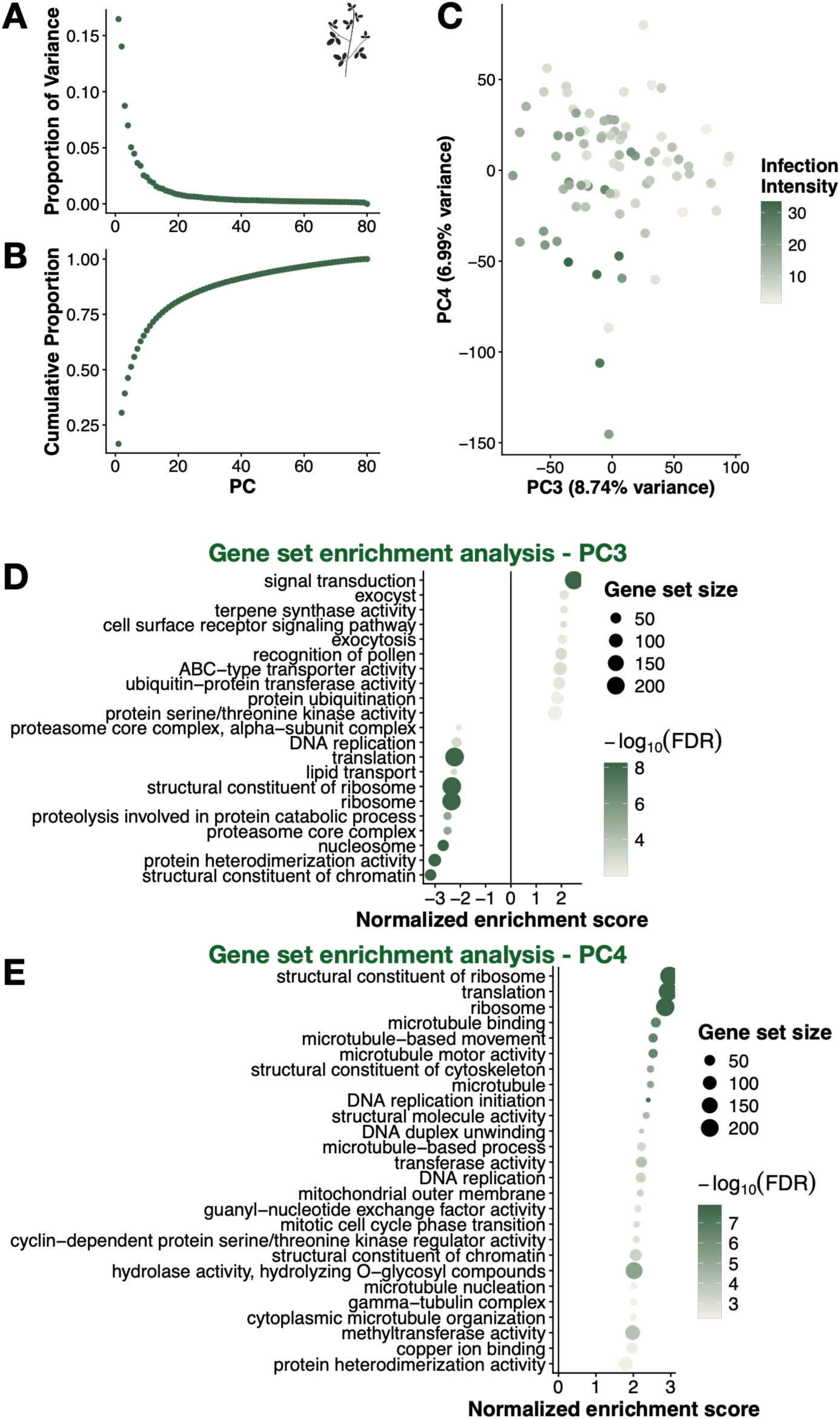
Host interaction gradients capture infection-associated developmental transcriptional programs. **A-B.** Variance explained by each principal component (PC) for the transcriptome of the plant host *Medicago truncatula*. **C.** PC analysis, showing host samples projected onto PCs 3 and 4, colored by infection intensity. PCs 3 and 4 collectively separate samples based on infection intensity. **D-E.** Gene ontology enrichment analysis of 300 genes with the most strongly positive and negative loadings into PCs 3 and 4. Both are enriched for biological processes associated with cell proliferation and growth, suggesting that infection intensity is associated with variation in host developmental activity during gall formation.

**Figure S5.**
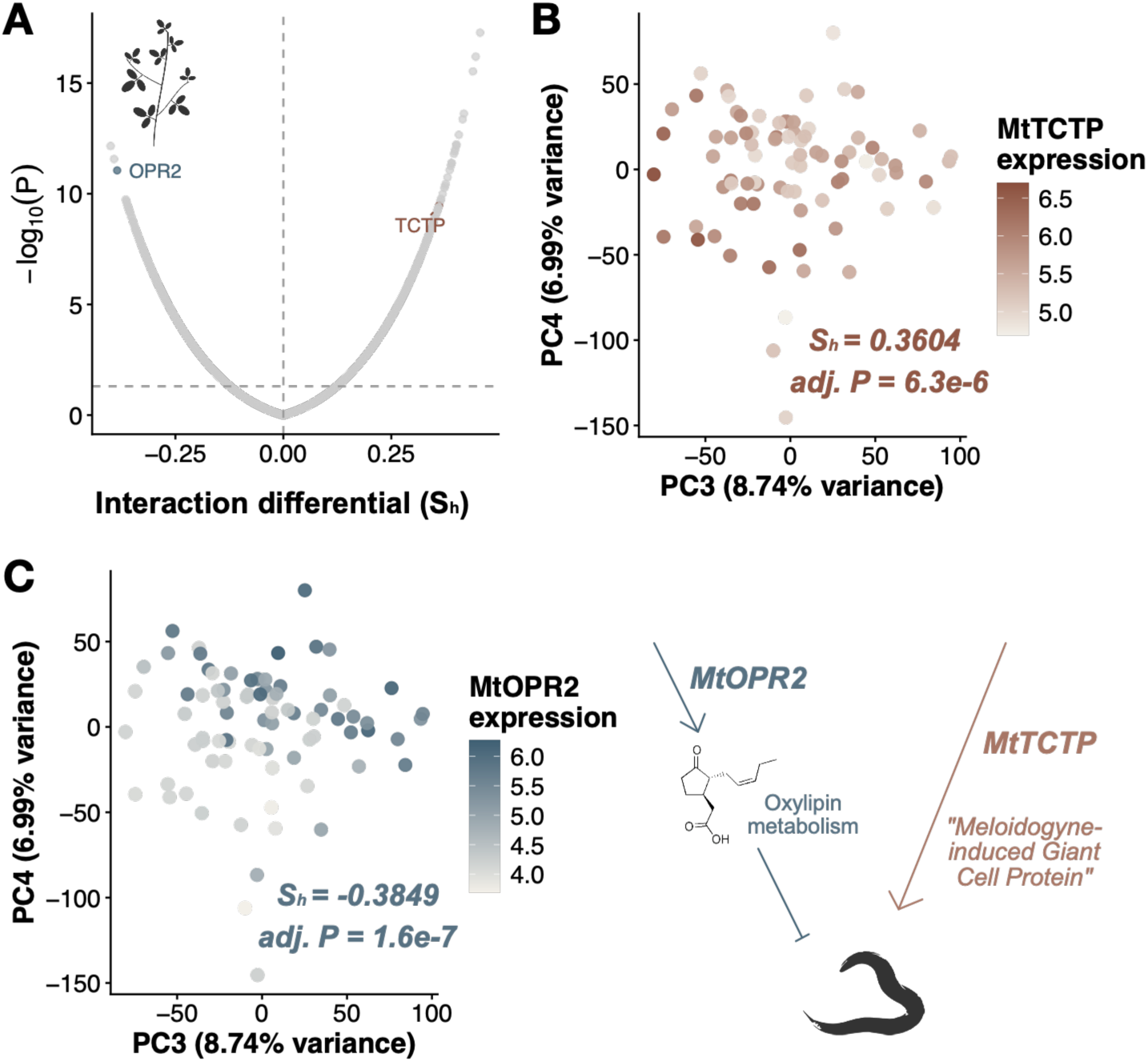
Interaction differentials identify known host genes associated with quantitative variation in infection outcomes. **A.** Volcano plot of interaction differentials (*S_h_*) for the plant host *Medicago truncatula* showing the relationship between effect size and statistical significance. *MtTCTP* (*TRANSLATIONALLY CONTROLLED TUMOR PROTEIN/MELOIDOGYNE-INDUCED GIANT CELL PROTEIN, MtrunA17_Chr1g0192201*), a feeding site-enriched transcript, and *MtOPR2* (*12-OXOPHYTODIENOATE (OPDA) REDUCTASE 2, MtrunA17_Chr5g0394891*), a gene involved in biosynthesis of oxylipins such as jasmonic acid, are highlighted as genes with strong positive and negative interactive differentials, respectively. **B.** Expression of *MtTCTP* shows positive covariance with infection intensity, consistent with a predicted function in feeding site establishment. **C.** Expression of *MtOPR2* displays negative covariance with infection intensity, consistent with positive contributions of *AtOPR2* to root-knot nematode resistance in *Arabidopsis thaliana* (Verhoeven et al. 2023).

**Figure S6.**
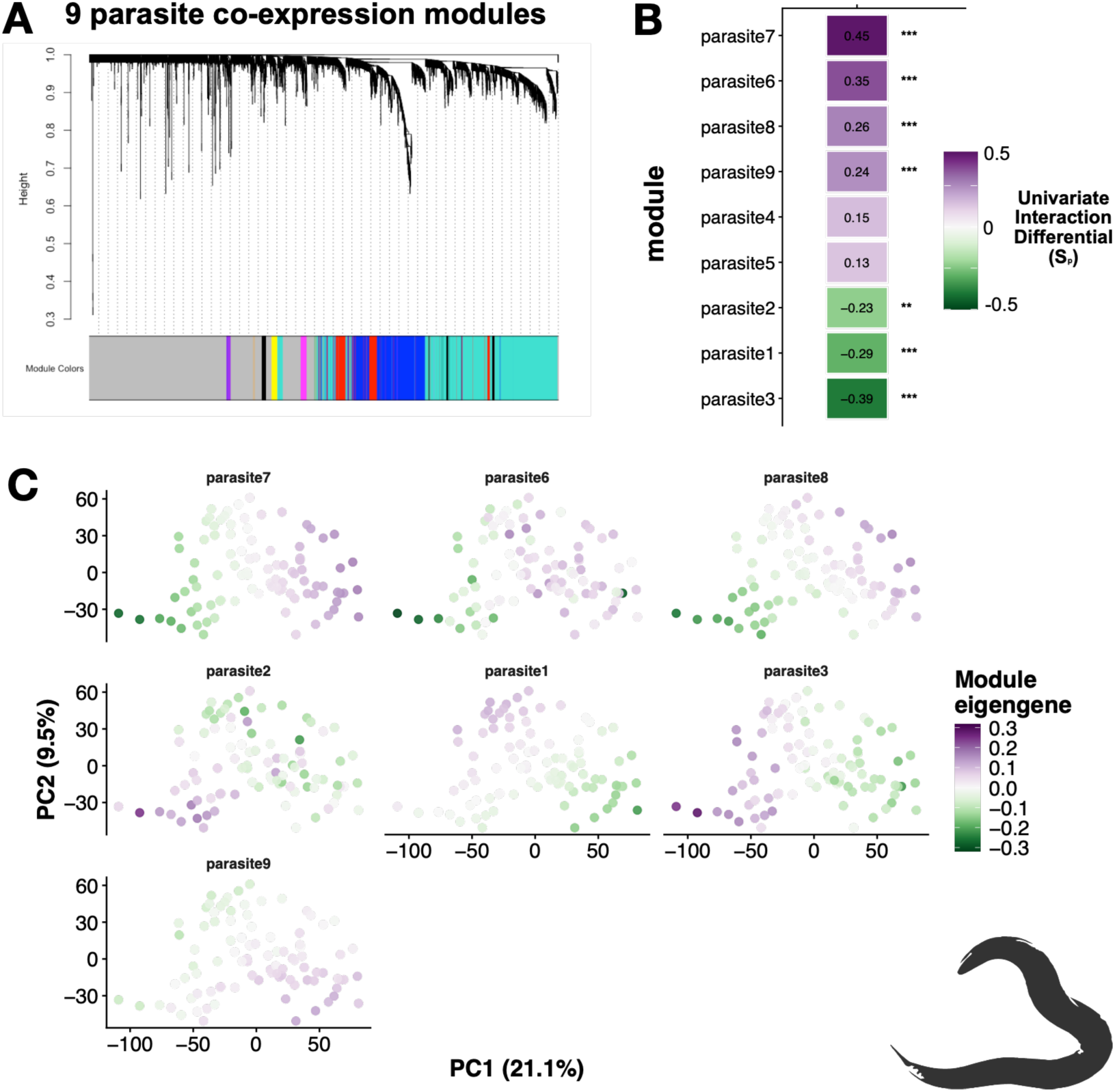
Dynamics of parasite transcriptional network modules along the interaction continuum. **A.** Hierarchical clustering of transcripts of the nematode parasite *Meloidogyne hapla* identified 9 co-expression modules using weighted gene co-expression network analysis (WGCNA). Gray indicates transcripts that were not assigned to a module. **B.** Interaction differentials (*S_p_*) estimated for each parasite module’s eigengene. Positive values indicate average module expression increases with infection intensity and vice versa. Asterisks denote Bonferroni-adjusted significance (*\*, adj. P < 0.05; **, adj. P < 0.01; ***, adj. P < 0.001*). **C.** Eigengenes for parasite modules with significant interaction differentials projected onto parasite principal component space. Modules associated with positive interaction differentials are generally associated with later parasite developmental stages, and vice versa.

**Figure S7.**
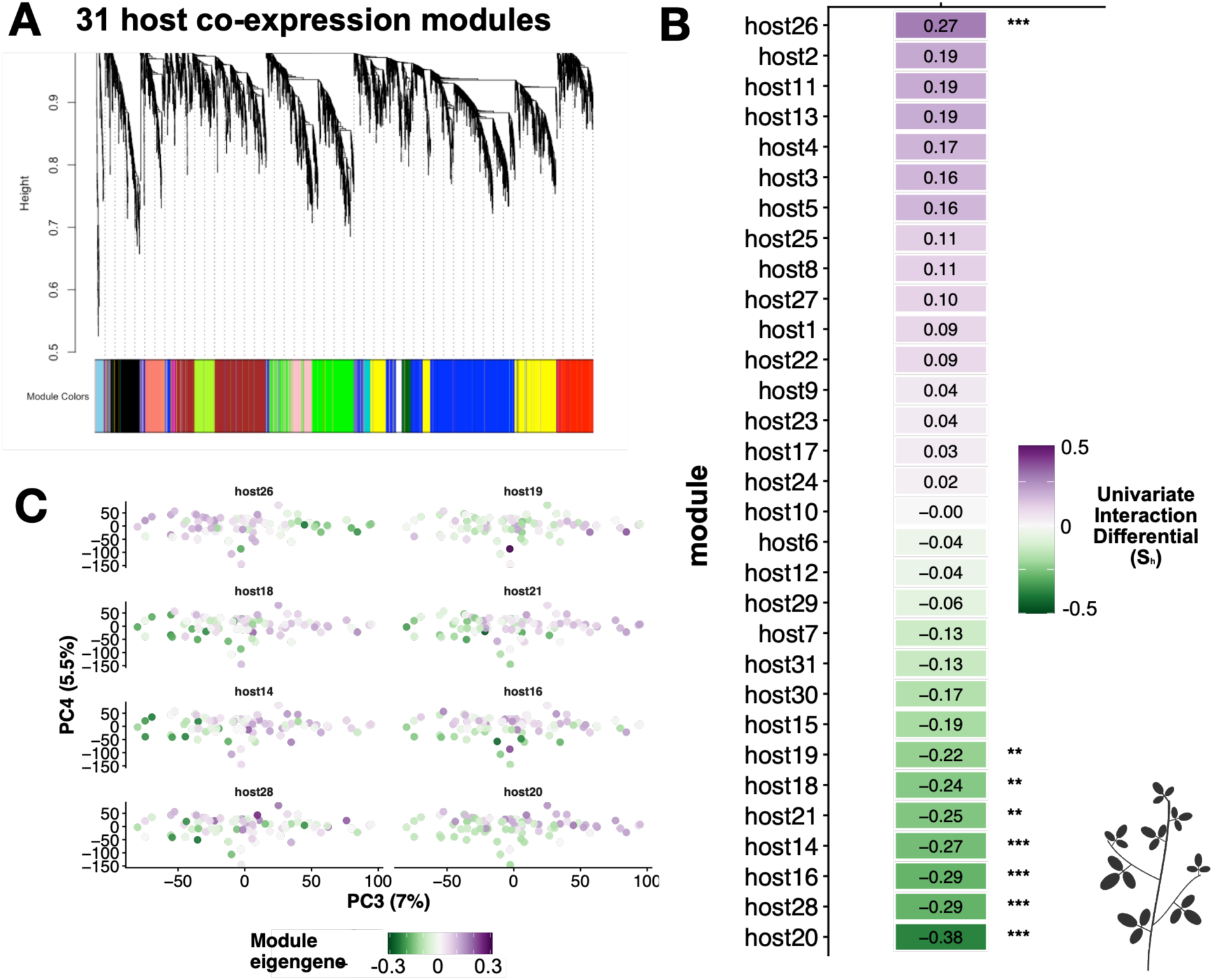
Dynamics of host transcriptional network modules along the interaction continuum. **A.** Hierarchical clustering of transcripts of the plant host *Medicago truncatula* identified 31 co-expression modules using weighted gene co-expression network analysis (WGCNA). **B.** Interaction differentials (*S_h_*) estimated for each host module’s eigengene. Positive values indicate average module expression increases with infection intensity and vice versa. Asterisks denote Bonferroni-adjusted significance (*\*, adj. P < 0.05; **, adj. P < 0.01; ***, adj. P < 0.001*). **C.** Eigengenes for host modules with significant interaction differentials projected onto host PCs 3 and 4 to illustrate their covariances with infection intensity.

**Figure S8.**
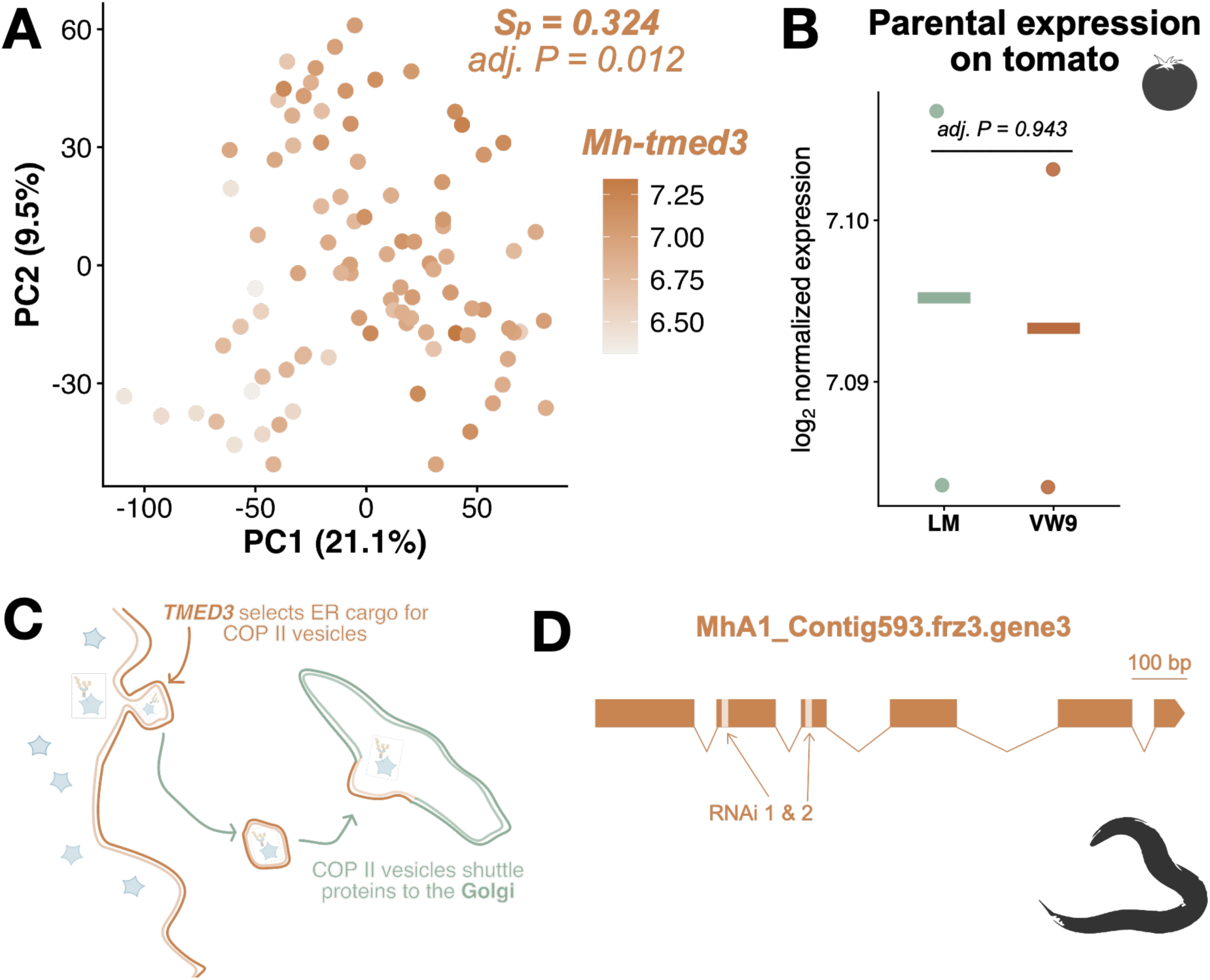
*TMED3* as a potential correlate of infection intensity. **A.** Abundance of *Mh-tmed3* transcripts across recombinant inbred lines (RILs) of the nematode parasite *Meloidogyne hapla* is highly correlated with infection intensity (*S_p_ = 0.324; adj. P = 0.012*). **B.** *Mh-tmed3* expression does not differ between parental *Mel. hapla* isolates VW9 and LM on tomato plants (*Solanum lycopersicum* cv. ‘Heinz 1706’; *adj. P = 0.943*). **C.** *Mh-tmed3* encodes an ERV25-related cargo receptor that facilitates incorporation of secretory proteins into COPII vesicles for transport from the endoplasmic reticulum to the Golgi apparatus, making it an essential component of the protein secretion machinery. **D.** Gene structure of *Mh-tmed3* showing regions targeted by dsRNA constructs used for RNAi.

**Figure S9.**
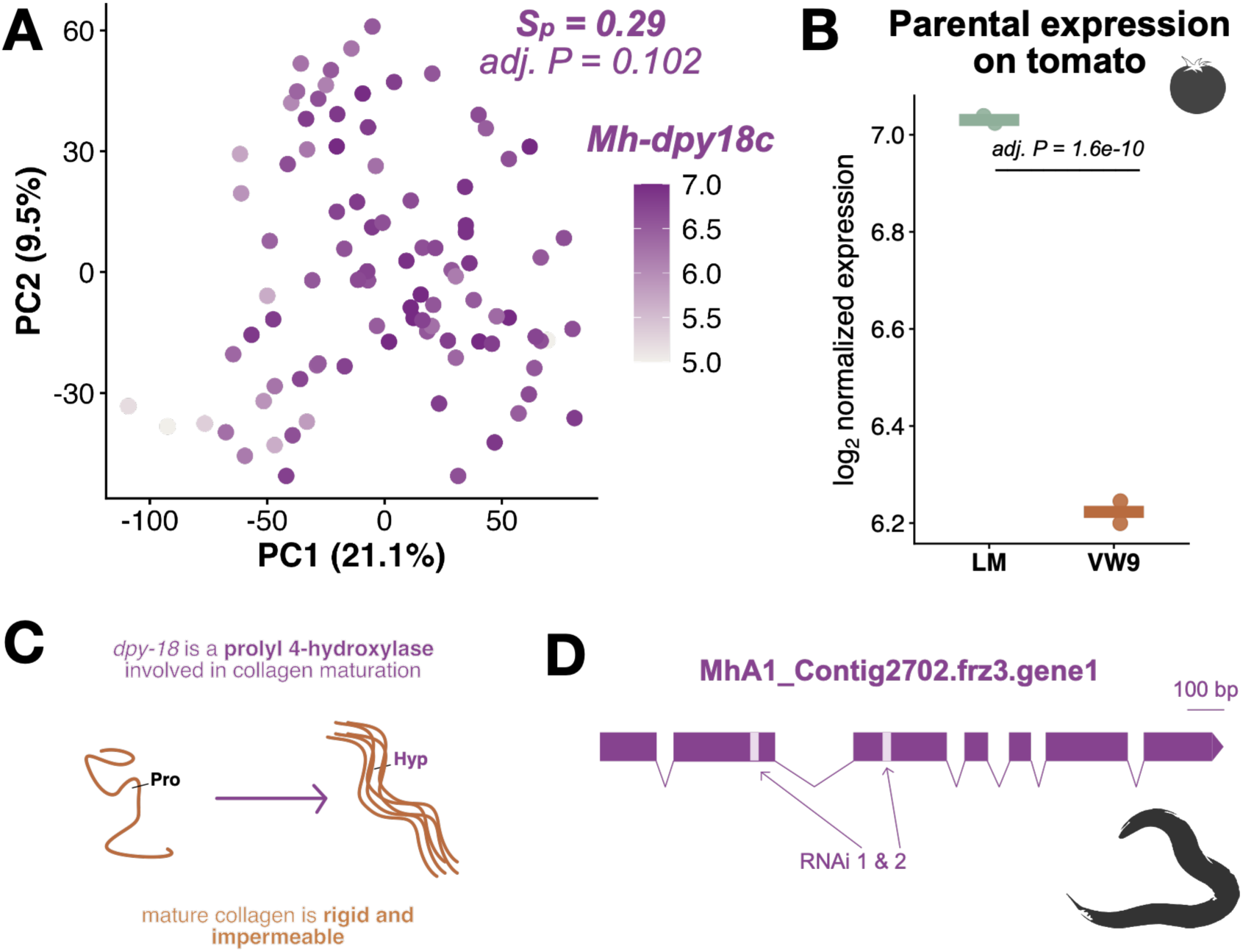
*DPY18c* as a potential correlate of infection intensity. **A.** Despite being a hub gene for the gene co-expression module ‘parasite6’ of the nematode parasite *Meloidogyne hapla*, abundance of *Mh-dpy18c* transcripts across parasite recombinant inbred lines (RILs) does not covary independently with infection intensity (*S_p_ = 0.29; adj. P = 0.102*). **B.** *Mh-dpy18c* expression does, however, differ between parental *Mel. hapla* isolates VW9 and LM on tomato plants (*Solanum lycopersicum* cv. ‘Heinz 1706’; *adj. P = 1.6×10^-10^*). **C.** *Mh-dpy18c* encodes a prolyl 4-hydroxylase that modifies procollagen peptides, enhancing their rigidity and impermeability. **D.** Gene structure of *Mh-dpy18c* showing regions targeted by dsRNA constructs used for RNAi.

**Figure S10.**
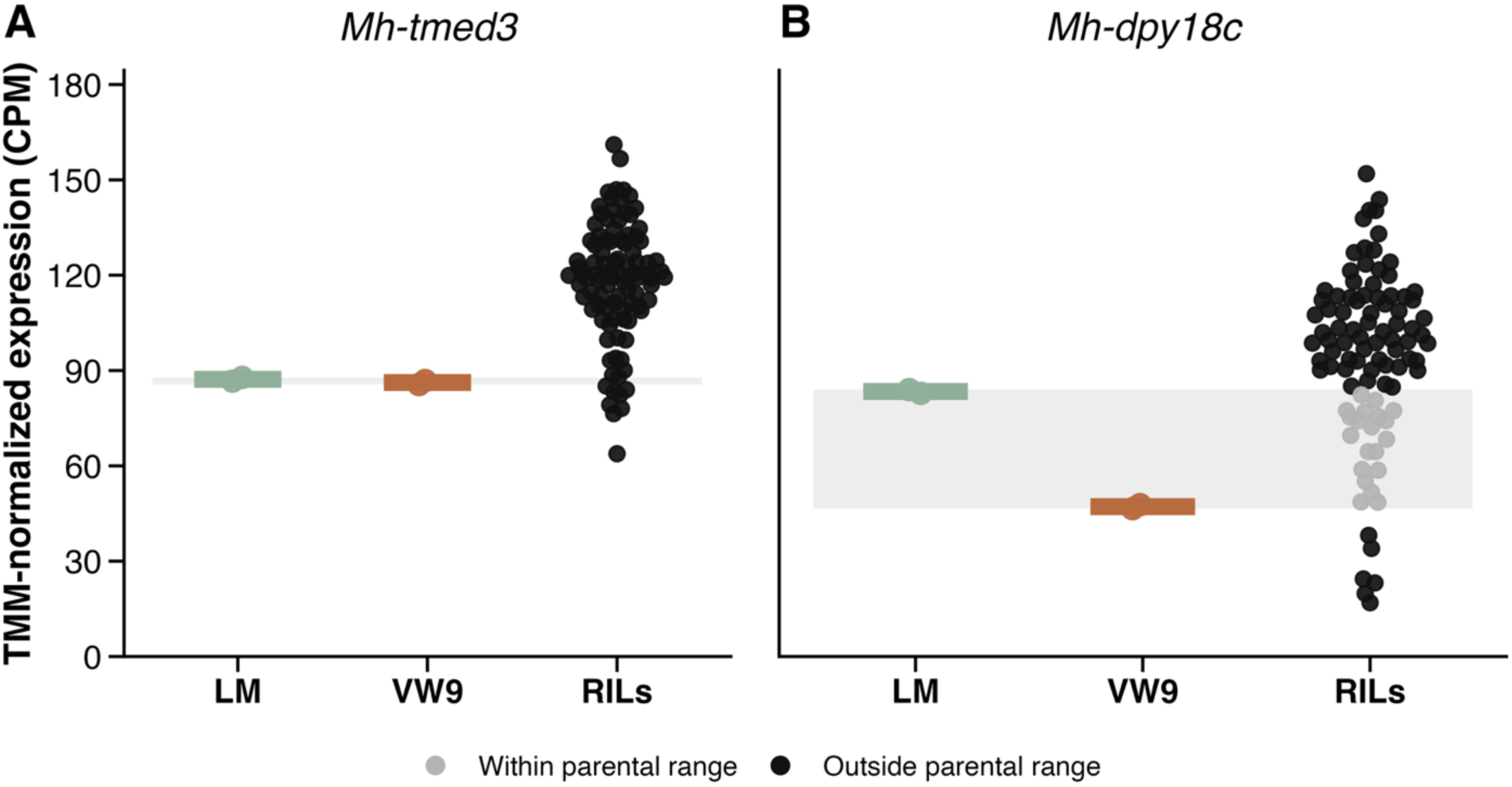
Mh-*tmed3* exhibits pronounced transgressive expression variation among recombinant inbred lines. TMM-normalized transcript abundance is shown for parental isolates LM and VW9 and 80 recombinant inbred lines (RILs). Horizontal bars indicate parental medians. Shaded area denotes the expression range observed across parental samples; RILs within this range are shown in grey, while RILs outside the parental range are shown in black and represent candidates for transgressive expression variation. (**A**) *Mh-tmed3* exhibited strongly directional variation (91.6% of RILs exceeded and 8% fell below the parental expression range. (**B**) *Mh-dpy18c* exhibited broader, bidirectional variation (72.9% RILs exceeded, 6.5% fell below).

